# Eyewire II – A connectomic resource for resolving cell types and circuits of the mouse retina

**DOI:** 10.64898/2026.05.28.727403

**Authors:** Sebastian Ströh, Simone Ebert, Julia Fadjukov, Jan Lause, Jonathan Oesterle, Katrin Franke, Ziwei Huang, Ran Lu, Arie Matsliah, David Berson, Venus Sherathiya, Dominic Gonschorek, Timm Schubert, Celia David, Marissa Sorek, Amy Sterling, Nikitas Serafetinidis, Joshua Singer, Yoshihiko Tsukamoto, Hiromi Maeyama, Naoko Omi, Gregory Schwartz, Philipp Berens, H. Sebastian Seung, Thomas Euler, The Eyewire II Consortium

## Abstract

Comprehensive wiring diagrams from electron microscopy (EM) are a powerful tool to understand the inner workings of the brain. The retina is an easily accessible part of the brain that performs complex visual computations. Its thin, layered structure offers a unique opportunity to decipher neural cell types and map their connectivity. A major obstacle has been the limited size of existing retinal EM datasets, which could not resolve rare cell types and neurons with large dendritic arbors. Here, we describe Eyewire II, a large-scale EM dataset covering nearly 1 mm^2^ of the adult mouse retina – roughly 10-100 times larger than previous retinal EM volumes. Human proofreading of an automated reconstruction has so far yielded more than 8, 000 bipolar cells, 13, 000 amacrine cells, and 4, 000 retinal ganglion cells. Automated detection is complete for synaptic ribbons and in progress for conventional synapses. Prior to EM imaging, visual responses to diverse stimuli – including natural movies – were recorded in a subset of neurons using two-photon Ca^2+^ imaging, enabling direct alignment of morphological and functional cell type identity. To enable high-throughput, automatic cell typing, we devised a human-in-the-loop approach that combines deep learning with human expert annotations. As a proof-of-principle, we show that morphological features of reconstructed bipolar cells are sufficient to recover all 15 known bipolar cell types with regular, non-overlapping mosaics. Together, these data and tools establish Eyewire II as a shared resource for the field of retina research. Already now, more than 30 laboratories worldwide are contributing proofreading, expert annotations, and software tools, advancing Eyewire II towards a complete cell type catalog and synaptic wiring diagram of a mammalian retina.

## Introduction

The retina is a highly structured neuronal tissue with a simple blueprint (1): Bipolar cells (BCs) receive input from photoreceptors and relay their signals to ganglion cells, the retina’s output neurons, whose axons form the optic nerve. Two classes of predominantly inhibitory interneurons – horizontal cells (HCs) and amacrine cells (ACs) – shape the signal flow from photoreceptors via BCs to retinal ganglion cells (RGCs). This simplicity, however, is deceptive: Retinal neurons have been estimated to comprise around 130 neuron types in most mammals (2, 3). ACs and RGCs constitute at least 80% of these types, exhibiting remarkable morphological, functional, and transcriptomic diversity (4–8). Given this diversity, the number of possible circuit configurations is large. Studying these circuits individually has been successful (for example in the rod pathway; 9–14), but uncovering the logic of the whole retinal network requires a larger scale approach.

Accelerating this progress requires finalizing the parts list of the retina (i.e. identifying all the cell types) and tracing out the wiring diagram. To obtain such information at scale, connectomics based on large-scale volumetric electron microscopy (EM) data has emerged as a powerful approach (15–21). Recent successful applications of EM connectomics relied on high-throughput deep learning algorithms for tissue segmentation and synapse detection, efficient web-based visualization, and active research communities that annotate, proofread, and analyze the data together (22–24).

Three shortcomings of existing retinal EM datasets have limited progress towards identifying retinal circuits and their function. First, available EM datasets have been severely limited in size, with the largest datasets (21, 25, 28) covering around 0.05 to 0.1 mm^2^ (29) – only around 0.6% of the whole mouse retina. This has made it difficult to reconstruct cells with larger dendritic arbors, as even for well-known medium-sized cell types such as the sustained On alpha (sOn-a) RGC (30), only a few truncated examples were included in these datasets, and most wide-field amacrine cell types are so large that they only exist as fragments. This means many important circuit parts – large RGCs and wide-field ACs – are not well covered by these earlier connectomes. Second, most

EM datasets do not include any functional information about the reconstructed cells such as responses to light stimuli, and those that do (21, 26) have used very limited stimulus sets such as moving bars. Third, some previous datasets lack staining of intracellular structures (25, 27), making it impossible to conclusively identify synapses. Other comparably small datasets include this information (for an overview, see 31), but segmentation and proofreading had been such a barrier that only a tiny fraction of the cells had been reconstructed.

Together, these three limitations make clear that the field is currently lacking a key community resource: A retinal EM dataset that is large enough to resolve most cell types, and includes both synapses and responses to rich visual stimuli. Such a resource would allow the field to complete the retina’s parts list and bring us closer to deciphering its visual computations.

Here, we present Eyewire II, an EM reconstruction of an almost 1 mm^2^ patch of mouse retina with readily visible synaptic structures, combined with functional responses to a diverse set of visual stimuli recorded *ex vivo* with two-photon (2p) Ca^2+^ imaging prior to EM. We used deep learning-based tools to speed up and improve segmentation, substantially reducing the effort for proofreading. Most BCs and small-field ACs required little or no proofreading. Currently, we have proofread >8, 000 BCs, >13, 000 ACs and >4, 000 RGCs, making this the largest retina connectomics dataset available. Finally, we provide software tools to extract morphologies, contacts, and highlight synapses, as well as web-based visualization tools ensuring that the dataset and its annotations are readily accessible and can be continuously refined through community efforts.

As a proof of principle, we developed a “human-in-the-loop” approach that combines machine learning and domain expertise to develop a comprehensive classification of all retinal neuron types. We use it to show how the dataset makes it possible to identify complete mosaics of all previously described BC types of the mouse retina. We also show that Eyewire II resolves all other retinal cell classes at type level, suggesting that the dataset will substantially advance completion of the retina’s parts list. For some of these types, we also highlight the connection between anatomy and function, providing a path forward for an integrated understanding of retinal cell types. We demonstrate that synaptic ribbons can be efficiently identified automatically throughout the reconstructed volume, which will facilitate identification of circuits and complete wiring diagrams.

## Results

Eyewire II is a new ultrastructural dataset of an adult, wildtype (C57Bl/6J) mouse retina. It has several advantages over earlier datasets: First, the dataset comprises about 1mm^2^ from the retina’s temporo-ventral quadrant (Fig. 1A-C), 10-100 times larger retinal area than earlier mouse EM datasets (21, 25–27). This size enables the reconstruction of substantial parts of even the largest amacrine cells, which span hundreds of micrometers (32, 33) (Fig. 1A), as well as complete mosaics of large RGC types. Second, unlike the previous largest mouse dataset (Eyewire I; 25, 26), Eyewire II was stained to reveal synapses and other intracellular features. Therefore, we can obtain a connectome based on actual synaptic connections, while Eyewire I had to resort to contact area between touching cells as a proxy. Third, as in Eyewire I, the retina underwent two-photon (2p) Ca^2+^ imaging before processing for EM. Where Eyewire I used only a simple moving bar stimulus, we presented a rich stimulus set, including the chirp stimulus and natural movies, yielding a dataset with more complex light responses. This links morphological and functional cell types for almost 400 cells recorded in the ganglion cell layer (GCL). Fourth, the Eyewire II dataset is accompanied by a suite of software tools to extract morphologies and analyze connectivity as well as annotations such as for synaptic ribbons.

**Figure 1.**
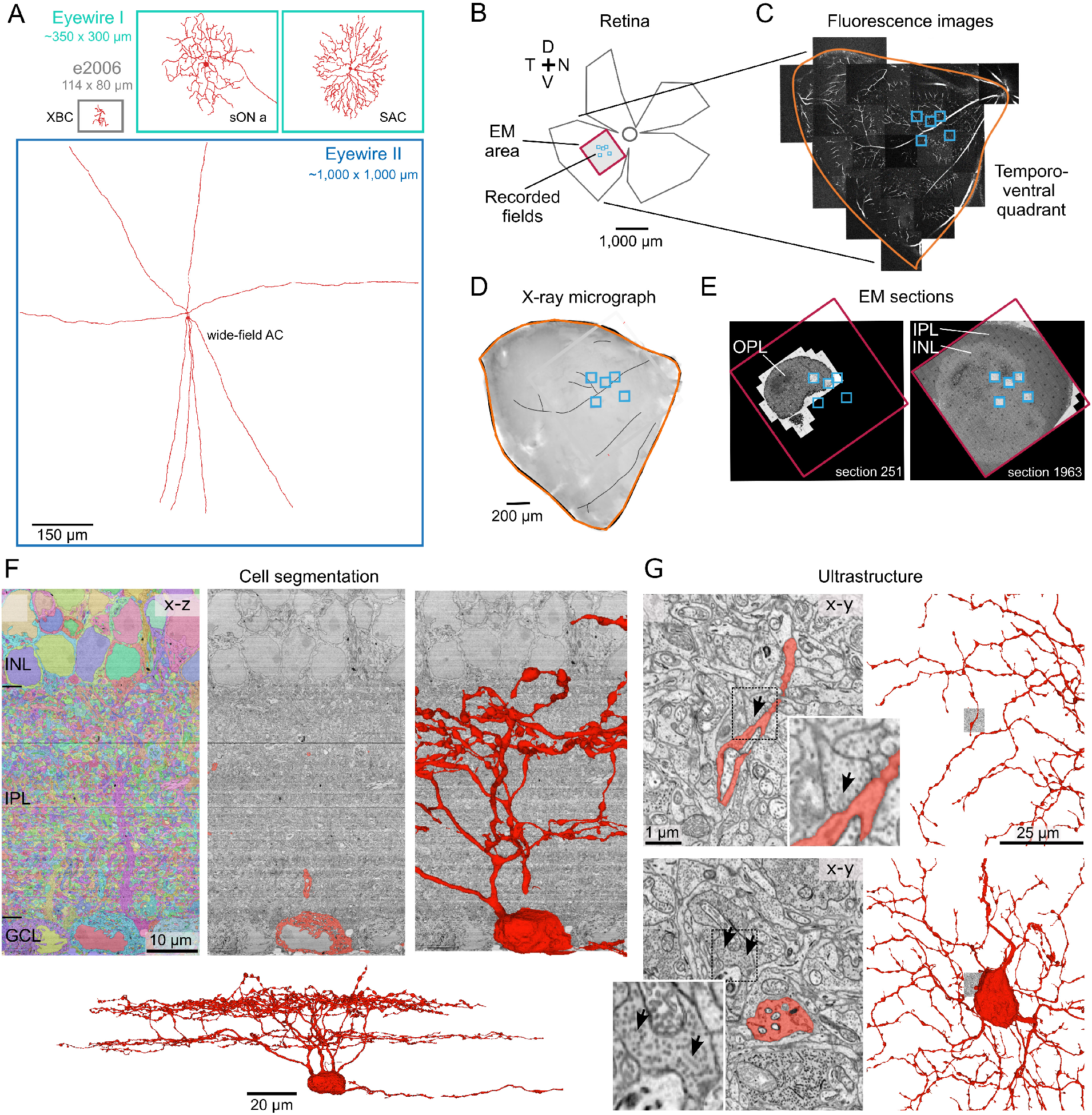
Dataset overview. (**A**) Eyewire II (dark blue) covers almost 1 mm^2^, comprising a much larger area than earlier EM datasets in mouse retina, such as Eyewire I (cyan; 25, 26) and e2006 (gray; 27), enabling reconstruction of substantial parts of even the largest wide-field amacrine cells (*bottom*).(**B**) Schematic outline of the imaged retina, with the positions of the five Ca^2+^ imaged fields (light blue) and the EM imaged area in the temporo-ventral retina (purple). (**C**,**D**) Temporo-ventral quadrant reconstructed from fluorescence images (Sulforhodamine 101, see Methods) taken directly after Ca^2+^ imaging in the recording chamber of the two-photon microscope (C) and as x-ray micrograph after fixation and embedding for EM (D). Orange outline shows approximate outline of quadrant. (**E**) Examples for EM sections, with outer plexiform layer (OPL) in small part of the image stack (*left*), and inner plexiform layer (IPL) and inner nuclear layer (INL) indicated (*right*). For movie of complete stack, see Fig. S1. (**F**) Vertical cross-section (x-z) through the current stack, with all segments color-coded (*left*), the segments of a retinal ganglion cell (RGC) labeled (red, *center*), its rendered morphology overlaid (*right*), and the complete RGC (*bottom*). (**G**) Examples for ultrastructure around the RGC’s dendrites, with examples for synaptic ribbons indicated (arrows; see insets).

### High-throughput, large-scale ultrastructural imaging of functionally characterized retina

To create the dataset, we loaded the tissue with a fluorescent sensor dye using electroporation and performed *ex-vivo* Ca^2+^ imaging of five 94*×*94 *µ*m^2^ fields in the temporo-ventral quadrant of an adult mouse retina using a standardized battery of visual stimuli, commonly used for functional cell type identification. This included a temporally modulated full-field “chirp” and a bright bar moving in 8 directions (34), as well as natural movies from mouse habitats (35). At the end of the recordings, we chemically fixed the tissue immediately and stained it for electron microscopy using heavy metals. Subsequently, we embedded the retina in epoxy resin for ultramicrotomy (Fig. 1D; see Methods).

We cut approximately 4, 000 40− 45-nm ultra-thin sections, of which 2, 064 have so far been imaged using a scanning EM with an imaging area of 1.05 × 1.05 mm (Fig. 1E) at a voxel size of 8 × 8 × 40 nm (segmentation and analysis at 16 × 16 nm). The majority of the sections contained the GCL with the somata of RGCs and displaced ACs, and the inner plexiform layer (IPL), where BCs, ACs, and RGCs form their synaptic connections. Many GCL and IPL sections cover around 1 mm^2^ of imaged tissue area, although tissue coverage somewhat decreased towards the inner nuclear layer (INL; Fig. 1E,F), where HC, BC, and regular AC somata are located. Some regions of the dataset include small portions of the outer plexiform layer (OPL; Fig. 1E), where BCs and HCs contact photoreceptors. Future imaging will extend the dataset to complete coverage of the OPL. Unlike in Eyewire I (25, 26), the tissue was processed to retain intracellular ultrastructure, enabling the detection and study of organelles such as synaptic ribbons, vesicles of conventional synapses, and mitochondria Fig. 1G).

### Segmentation and proofreading

After alignment of the sections, we automatically segmented the ultrastructure using deep learning algorithms (see Methods), resulting in almost complete cells (“AI meshes”, like the cell in Fig. 1F-G; see also Fig. 3A). Typical segmentation errors included the merging of two unrelated structures and incomplete dendritic arbors (Fig. 2A,B). Many of these errors could be easily spotted by humans and fixed, e.g., the meshes of a BC and a AC being merged in one point or a large gap in a dendritic arbor of a RGC – and could be easily fixed during proofreading.

**Figure 2.**
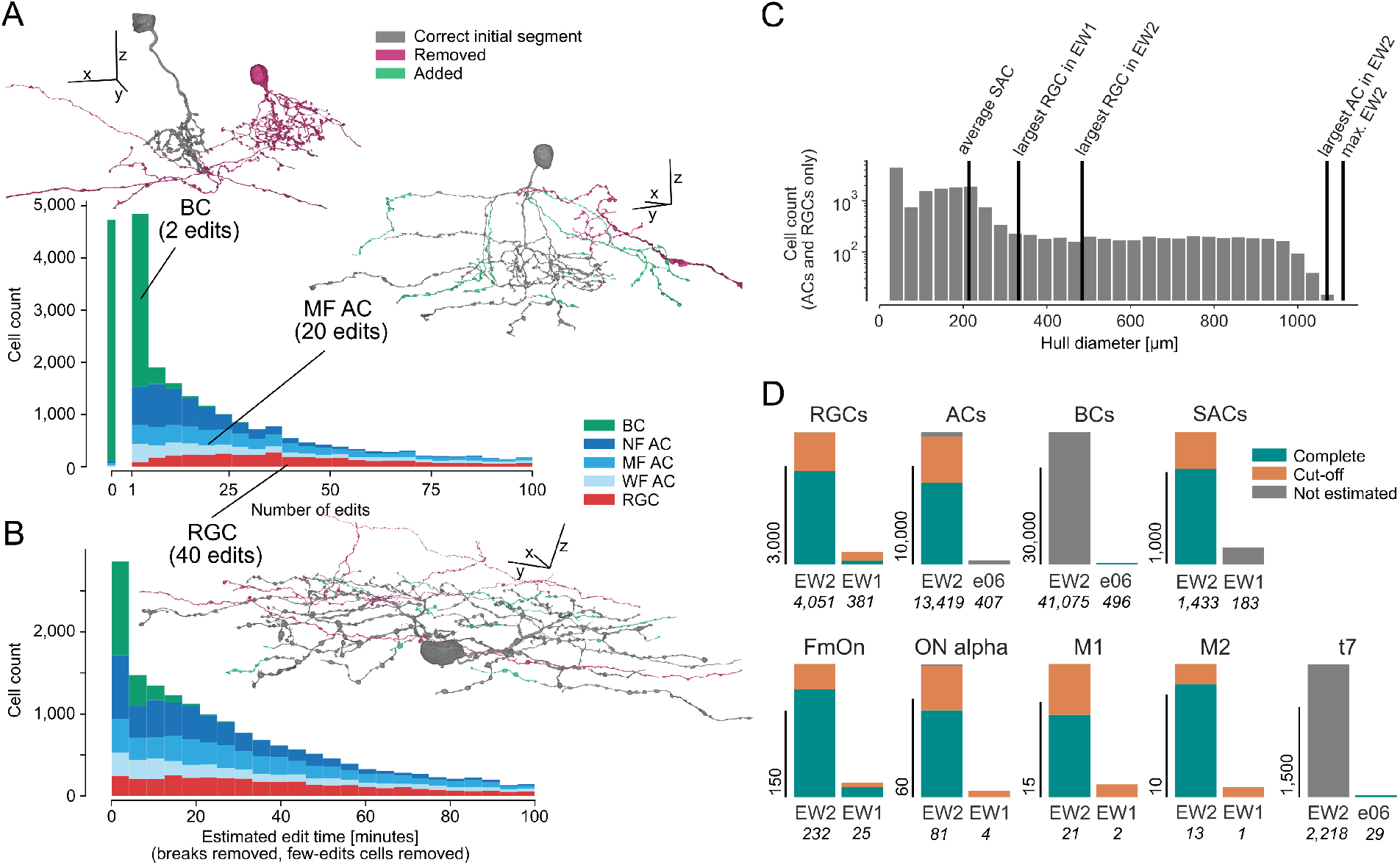
Proofreading statistics. (**A**) Examples for typical edits from each of the major cell classes, with the initial correct segment (gray) as well as removed (magenta) and added (green) segments indicated. The histogram shows the number of edits needed per proofread RGC (red), AC (blue), and BC (green). The bar at 0 reflects cells that did not need any edits. (**B**) Distribution of estimated edit time per proofread cell (colors as in (A)). Cells with less than two edits and proofreading breaks longer than 15 minutes were ignored. (**C**) Circle-equivalent diameters from convex hulls of RGCs (ignoring RGC axons) and ACs with reference values. (**D**) Cell counts sorted by either cell class or cell type. Reconstruction status, i.e., if cells were reconstructed completely or if they are cutoff at the volume borders, is color-coded (light-blue, complete; orange, cut-off) or gray if this information was not available. Comparison against Eyewire I (EW1; 22) or e2006 (e06; 27).

**Figure 3.**
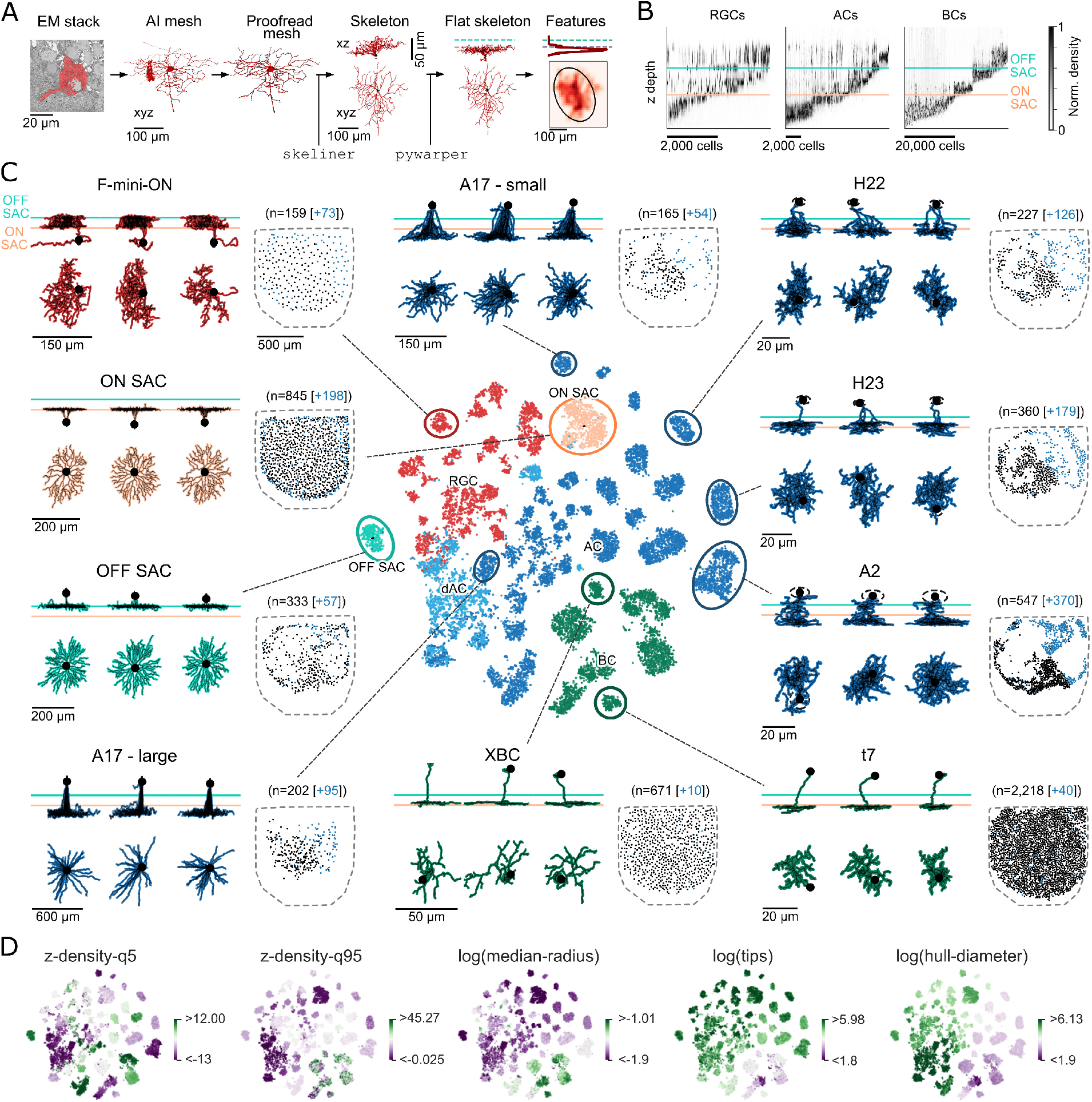
Large-scale dataset enables comprehensive morphological classification of retinal neurons. (**A**) Illustration of the processing pipeline. From left to right: The EM stack was segmented using a previously developed algorithm. The automatically segmented cell meshes were proofread by professional proofreaders and researchers. We computed skeletons from the meshes using `skeliner`, which were then flattened using `pywarper`. From the flattened skeletons, we computed morphological features, such as density profiles. (**B**) Normalized densities along z-axis across the inner retina for all 59, 000 neurons split by cell class: retinal ganglion cells (RGCs), amacrine cells (ACs), and bipolar cells (BCs). Cyan and apricot-colored lines indicate stratification levels of Off and On starburst cells (SACs), respectively. (**C**) t-SNE computed from 43 normalized morphological features of all proofread RGCs (red), ACs (blue; except On SACs, apricot, and Off SACs, cyan), and a subset of 200 BCs per type (green), with representative example neurons in x-z and x-y view, as well as cell distribution (dashed outline, outline of IPL; manually annotated cells, black dots; additionally automatically classified cells, blue dots). (**D**) Distributions of selected features (for details, see Methods).

Due to the high quality of the initial segmentation, proofreading was fast: The median number of required edits per cell ranged from 0 for BCs and 14 for narrow-field ACs (NF ACs) to 33 for medium-field ACs (MF ACs), 35 for wide-field ACs (WF ACs), and 41 for RGCs (Fig. 2A), equivalent to median edit times of 2 (BCs), 19 (NF ACs), 34 (MF ACs), 39 (RGCs), and 41 (WF ACs) minutes per cell (Fig. 2B). At the time of writing, >25, 000 of the estimated 100, 000 BCs, ACs and RGCs in the volume had been proofread by a group of 28 researchers, 72 paid and trained specialists, and 11 volunteer citizen scientists over the course of 12 months. These proofread cells included >4, 000 RGCs (90% of all expected RGCs), >13, 000 ACs (29%), and >8, 000 BCs (16%), with complete coverage of a central 700 × 700 *µm*^2^ square in the GCL. Considering that proofreading so far focused on complex RGCs (which require many more edits than e.g., BCs and NF ACs), we estimate that with 25% of all expected cells proofread, we have already completed 35% of the total proof-reading effort required. The proofread cells in Eyewire II include hundreds of large cells, from the largest RGC types to most wide-field ACs (Fig. 2C), and lack only the likely rare, super-wide-field ACs that span more than 1 mm (e.g., 39). Already at the current stage of proofreading, Eyewire II provides 5-10 times more cells per type than previous datasets (Fig. 2D).

### Human-in-the-loop approach to morphological typing of retinal neurons

After proofreading, we extracted and skeletonized cell morphologies (Fig. 3A; for details on the pipeline, see Methods). Since the retinal tissue was slightly curved, we locally corrected the z-profile of cells using the dendritic plexuses of a subset of overlapping On and Off starburst ACs (SACs) as landmarks (40). This allowed us to precisely relate morphological features, such as dendritic arbor density (Fig. 3B), to normalized IPL depth.

From the flattened skeletons, we extracted morphological features including density profiles and summary statistics, such as the number of branch points or hull diameter. To explore this feature space, we visualized the reconstructed cells in a two-dimensional embedding (Fig. 3C,D; for a complete list of morphological features used, see Methods). We found that many well-known cell types separated into distinct islands, indicating a rich feature space that resolved biologically relevant morphological variation between retinal cell types.

To make Eyewire II useful to the community, researchers need to be able to immediately find their cells of interest, typically by looking up the cell type of proofread cells. At the same time, the large number of cells in the dataset precludes manual cell type annotation. This is why we used the morphological features described above to automatically identify the cell type of proofread cells – to enable cell type related analysis at scale. Here, we chose a “human-in-the-loop” procedure to obtain cell type labels efficiently, combining the scalability of machine learning approaches with the expertise and literature knowledge of domain experts (for details, see Methods). Our goal was not a definitive, automatic classification of all retinal cell types, although we demonstrate the feasibility of such an approach for BCs below (see Automatic BC type classification).

In brief, human expert annotators suggested initial cell class and cell type labels for 3D morphologies. These labels were then used to train random forest classifiers to make predictions for both already labeled, but also for previously unlabeled cells based on the automatically extracted features. Cells with a mismatch between human and classifier label were revisited by human experts. Once the dataset grew, we were able to employ unsupervised clustering to propose cell clusters of morphologically similar cells to speed up the process of cell typing even further. To this end, we mapped cell clusters to existing cell labels. Using this approach, we were able to identify most known retinal cell types (Fig. 3C; see also Fig. 4, Fig. 5). The cell class and type labels reported here are a consensus between export labels and classifier output (for details, see Methods).

**Figure 4.**
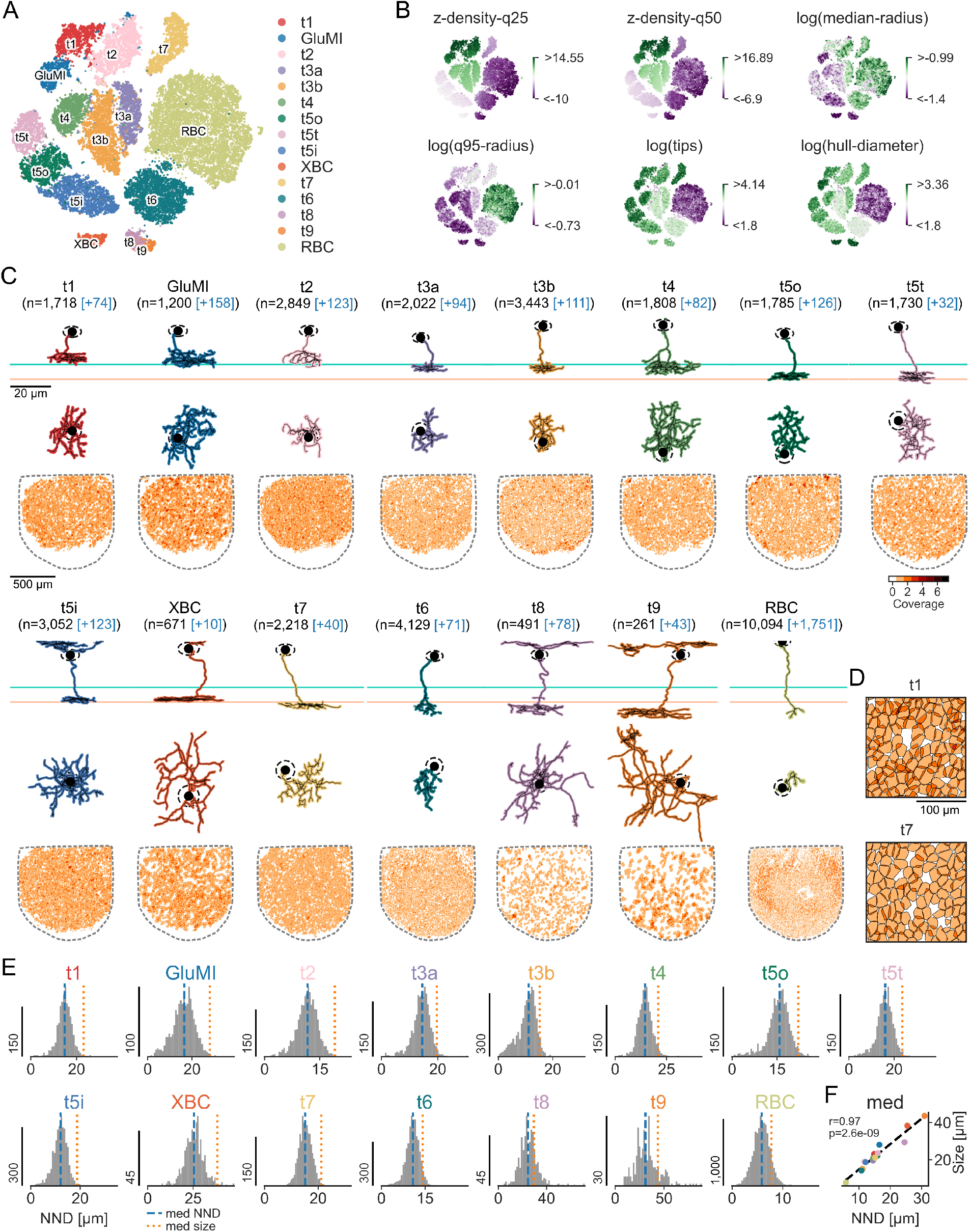
Automated clustering of bipolar cells based on axonal morphology and IPL depth. (**A**) t-SNE computed from 21 normalized axonal morphological features of 41, 000 bipolar cells (BCs). Colors indicate known BC types (27, 36–38). 700 BCs with no type label are shown in gray. Note that t8 and t9 are more difficult to separate based on the used features. (**B**) Distributions of selected features (for details, see Methods). (**C**) All BC types from (A), each with representative example in x-z (*top*) and x-y view (*center*) sorted by axonal IPL depth, and mosaic of axon terminals in x-y view (*bottom*). Colors as in (A); cyan and apricot-colored lines indicate stratification levels of Off and On starburst cells (SACs), respectively. (**D**) Zoomed-in mosaics of t1 and t7 BCs. (**E**) Nearest-neighbor distance (NND) for each BC type. (**F**) Axon terminal size (circle equivalent diameter) as a function of NND.

**Figure 5.**
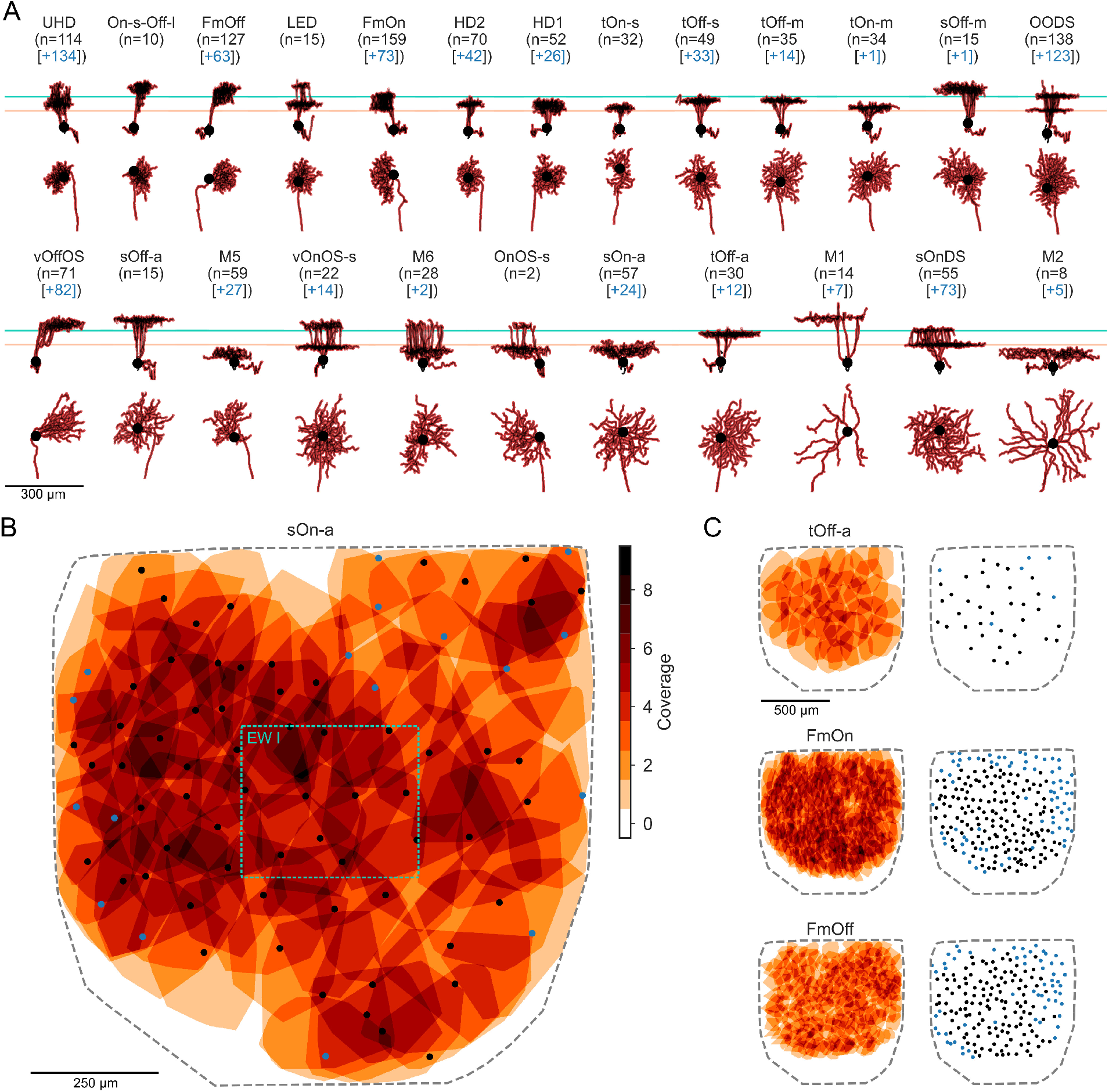
Morphological retinal ganglion cell (RGC) types. (**A**) Representative example cells for expert-annotated types that have been described previously (e.g., 22, 45, 46). Each type is shown in x-z (top) and x-y view (bottom). From top left to bottom right, types are sorted by dendritic arbor size. Horizontal lines indicate dendritic plexuses of Off SACs (cyan) and On SACs (apricot). *UHD*, ultra high definition; *On-s-Off-l*, On small Off large; *FmOff*, F-mini Off; *LED*, local edge detector; *FmOn*, F-mini On; *HD2*, high definition 2; *HD1*, high definition 1; *tOn-s*, On transient small RF; *tOff-s*, transient Off small; *tOff-m*, transient Off medium; *tOn-m*, On transient medium RF; *sOff-m*, Off medium sustained; *OODS*, On-Off DS; *vOffOS*, Off vertical OS; *sOff-a*, sustained Off *α*; *M5*, melanopsin 5; *vOnOS-s*, On vertical OS small RF; *M6*, melanopsin 6; *OnOS-s*, On OS small RF; *sOn-a*, sustained On *α*; *tOff-a*, transient Off *α*; *M1*, melanopsin 1; *sOnDS*, On DS sustained. *M2*, melanopsin 2; (**B**,**C**) Mosaics of the dendritic arbors of selected RGC types (x-y view) plus soma positions (expert annotation, black dots; classifier prediction, blue dots). We excluded 6 cells here, because the automatic axon detection failed for them, resulting in incorrect dendritic arbor convex hulls. Area of Eyewire I indicated as green box in (B).

Clusters were flagged as potentially novel cell types if they were not dominated by any known cell type or if known cell type labels split into clear subclusters. For example, the A17 amacrine cell – a well-characterized interneuron of the rod pathway (41, 42) – separated into two distinct clusters differing primarily in dendritic arbor size (Fig. 3C, *top-center* vs *bottom-left*). Whether these represent genuine types with distinct circuit roles, or morphological variation within a single type, remains to be determined. This illustrates how the combination of large cell numbers and systematic clustering can reveal fine-grained distinctions that are difficult to detect in smaller datasets.

### Automatic BC type classification

We applied this approach to BCs, combining it with automated synaptic ribbon detection (see Methods) to provide a classifier for BCs (Fig. 4A). The synaptic ribbon detector automatically identified 42, 000 BC candidates with > 35 ribbons (for details see Methods). Using morphological features of the BC axon terminals, such as density in the IPL and hull diameter (for a selection of features, see Fig. 4B), we recovered all 15 previously described types (27, 36–38) and their mosaics (Fig. 4C,D). To quantitatively test mosaic quality, we computed nearest-neighbor distance (NND) between neighboring cells of the same type (Fig. 4E), indicating very few mosaic violations. Furthermore, we found that NND correlated strongly with axon terminal size (*r* = 0.97, *p* < 10^*−*8^, *n* = 15; Fig. 4F), suggesting regular mosaics following similar scaling laws. This classification included 95% of all BC candidates. We assume that most of the remaining cells will belong to these 15 BC types, filling the few remaining gaps in the mosaic. It remains possible that some of these cells are non-canonical BC types (e.g., 43, 44), warranting further exploration in future work.

### Relating morphological RGC types to Ca^2+^ responses

We identified examples of almost all well-known RGC types (Fig. 5A) and obtained large, well-structured mosaics for many of them (Fig. 5B,C), including large RGCs, such as the alpha types. This allowed us to relate morphological RGC types to their light responses obtained prior to EM imaging (see Methods; Fig. 6A-C). To this end, we used high-resolution images of each recording field to register the recorded GCL cell somata (*n* =224 RGCs, *n* =153 displaced ACs) with the segmented ones (e.g., Fig. 6C vs. D). This enabled us to align morphology (Fig. 6E,F) and Ca^2+^ responses (Fig. 6G) for every soma in the recorded fields.

**Figure 6.**
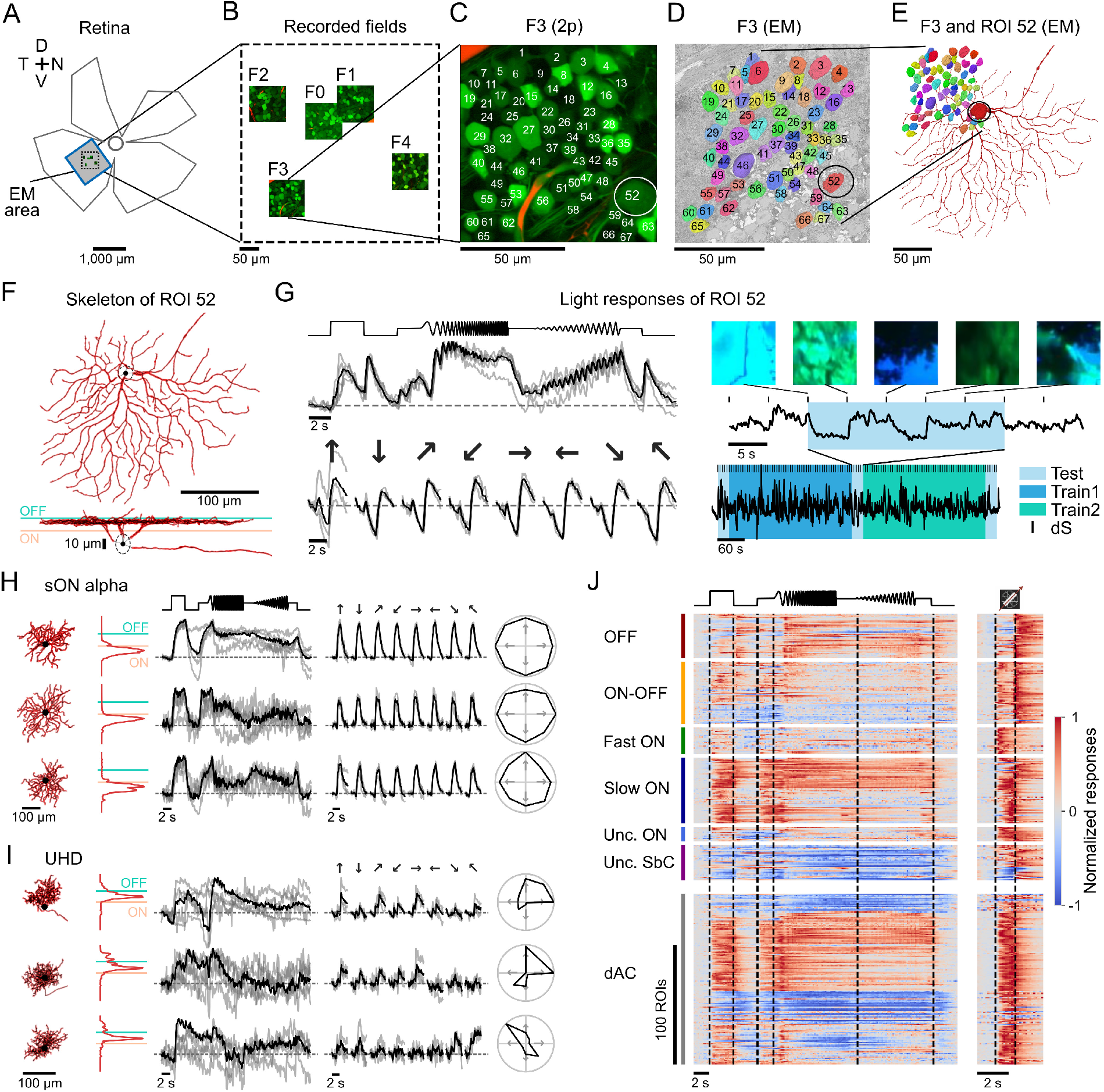
Overview of Ca2+ recordings. (**A**) Retinal outline with the five recording fields (dashed black box; cf. (B)) and the EM imaged area (gray box) located in the temporo-ventral retina. (**B**) The five fields F0 to F4 from (A), enlarged. (**C**) Recording field F3 from two-photon (2p) Ca^2+^ imaging, with ganglion cell layer (GCL) somata labeled by fluorescent indicator Oregon-Green BAPTA-1 (green); numbers indicate ROIs of recorded cells. (**D**) F3 mapped to the EM stack. Nuclei of cells corresponding to ROIs in two-photon (2p) recordings are shown in color; the nucleus of GCL cell corresponding to ROI 52 (circle in (C)) is highlighted. (**E**) As in (D), but showing the complete morphology (“mesh”) of cell corresponding to ROI 52. (**F**) Reconstructed skeleton of same cell as in (E) in top and side view (cyan and apricot lines indicate Off and On starburst amacrine (SAC) cell stratification bands, respectively). (**G**) Normalized Ca^2+^ responses for the chirp (*top-left*) and moving bar (*bottom-left*) – shown as trials (gray) and average (black) – to the natural movie stimuli (*right*), with example frames (*top*) at clip transitions (dS, black ticks), zoomed-in (*center*), and whole trace (*bottom*). Stimulus sections (Test, Train 1 and 2) are marked by colored boxes. (**H**,**I**) Examples for selected retinal ganglion cell (RGC) types, identified by morphology. In each panel, *from left to right:* Morphology from EM (x-y view, IPL stratification profile relative to On and Off SACs in cyan, and apricot, respectively), Ca^2+^ responses to chirp and moving bar, and directional tuning curve (Ca^2+^ responses as trials, gray, and average, black). (**J**) Normalized responses for chirp stimulus (*left*, as averages over repetitions) and moving bar (*right*, as projections to the preferred direction), split into 224 RGCs and 153 displaced amacrine cells (dACs) based on morphology, and then sorted by functional super-groups (34).

For many morphological RGC types, we found consistent responses, such as for sON-a cells (30, 47) (Fig. 6H). Some RGC types, however, showed variations in their light responses that can be partially explained by dendritic arbor size and position relative to the stimulus, which was centered on the recording field. For example, the small UHD cells, which have strong surround suppression (48), displayed less consistent responses, especially to the moving bar (Fig. 6I). Among all imaged cells, the functional diversity matched that previously observed both for RGCs and ACs (Fig. 6J).

### Responses change across recording fields

For some cell types, light responses differed systematically across the five recording fields (F0 to F4), reminiscent of response changes observed in an earlier study (49). On SACs were among the cell types showing the most dramatic response changes (Fig. 7). For the chirp stimulus, they displayed delayed On responses to the step and the beginning of the frequency chirp in F0, but then became increasingly suppressed in F1 to F3, before recovering in F4 (Fig. 7A,B). Moreover, the moving bar responses became increasingly delayed from F1 to F3, before loosing the delay again in F4 (Fig. 7B, *bottom* row).

**Figure 7.**
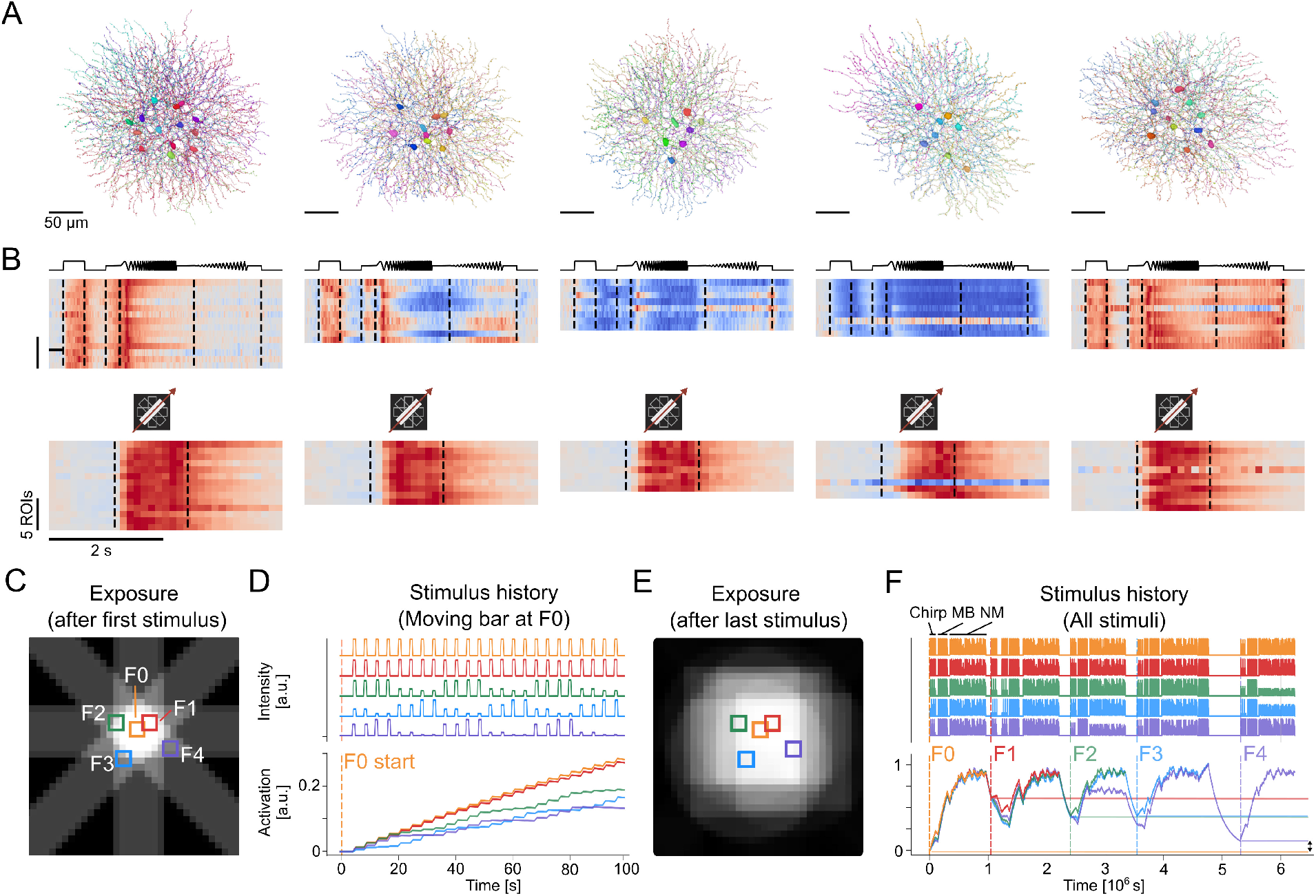
Starburst amacrine cell (SAC) responses change dependent on light exposure. (**A**) Reconstructions of morphologically identified On SACs in the 5 recording fields with Ca^2+^ responses. (**B**) SAC Ca^2+^ responses by field for chirp (*top*) and moving bar (*bottom*) stimuli as heatmaps with each line representing a cell. (**C**,**D**) Light exposure of the 5 fields until the end of the first stimulus presentation (moving bar centered on F0) visualized as map (C) and as stimulus intensity (D, *top*) and as activation over time by recording field (D, *bottom*. Activation was implemented as a leaky integrator with *τ* =240 s (for details, see Methods). (**E**,**F**) Like (C,D) but until after the last stimulus presentation (natural movie stimulus to F4). For clarity, activation traces are only depicted until the end of the respective field’s recording. Horizontal lines indicate activation level at the start of each recording field; as indicated by the arrow, initial activation for F0 and F4 are closer to each other than to those of the other fields..

We hypothesized that differences in light adaptation contributed to the observed response variability, because the fields were spatially close to each other such that later recorded fields had been exposed to stimuli presented to earlier recorded fields. To test this hypothesis, we calculated the light exposure of the recording fields during the 2p imaging experiment ((Fig. 7C,E; see Methods). Based on stimulus intensity trace (Fig. 7D,F, *top*), we estimated activation (*bottom*) separately for each field using a leaky integrator (with *τ* = 240 s) to account for light adaptation.

We found that activation at the start of the recording differed between fields, as cells in F4 had more time to recover from the previous stimulus presentations than F1, F2 and F3 (due to a longer pause between F3 and F4; Fig. 7F). Because the chirp and natural movie stimuli were large compared to the relatively short distances between recording fields, stimulus location likely played no major role – except maybe for F4, which sits slightly more to the edge of the cumulated stimulus (Fig. 7E). This suggests that light adaptation may have contributed to the observed cell type specific response differences between fields.

#### Reconstructing synaptic circuits

Towards understanding how the diverse functional responses emerge in the retina, we explored whether retinal circuits could identified with the help of the clearly visible synapses (Fig. 8A-C). We manually annotated conventional and ribbon synapses in a sub-volume of the dataset spanning 14× 20 *µ*m across the entire IPL. We extracted all proofread and typed cells in which at least one synapse was detected. The resulting cell type-specific connectivity graph included *n* = 9 rod bipolar cells (RBCs), whose inputs and outputs were quantified (Fig. 8D). At least four AC types connected to RBCs (Fig. 8E): The RBCs mainly provided output to A17 (*n* = 45) and A2 (*n* = 4) ACs, which are known to be key elements of the rod pathway (9). They received input from A17s, nNOS-1s, and a medium-field AC type (Fig. 8D): A17 cells provided the majority of inhibitory inputs into RBCs, confirming the well-studied reciprocal connectivity between RBCs and A17 (50). We found *n* = 8 wide-field nNOS-1 ACs, which contribute inhibition to the surround responses in A2 ACs (51). Those cells also received sparse input from RBCs (Fig. 8D). The remaining *n* = 4 ACs were tri-stratified medium-field ACs and may correspond to type H42 or H52 (27). The latter cells could be the source of glycinergic input to RBCs, which has been previously reported (52).

**Figure 8.**
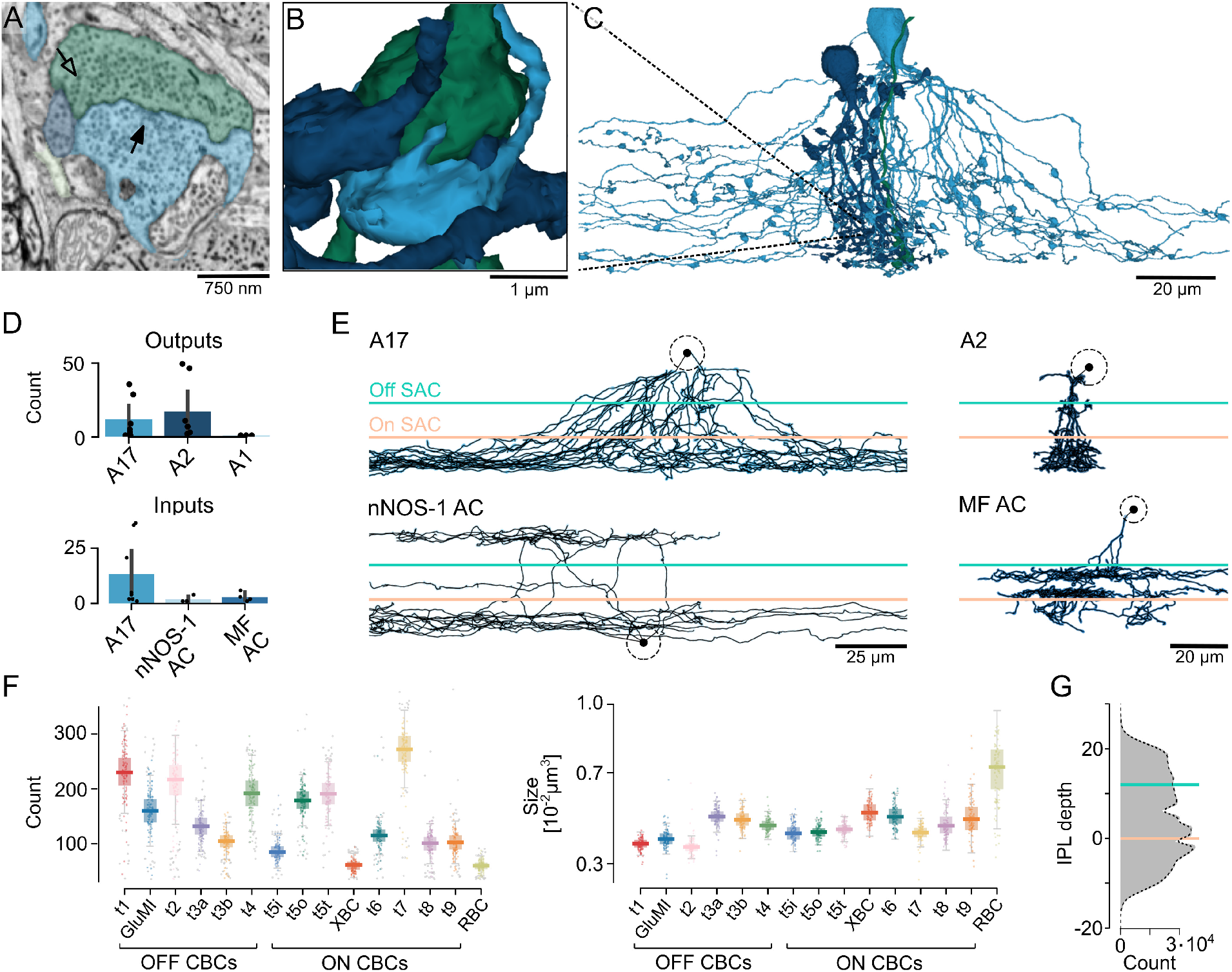
Towards large-scale analysis cell type specific synaptic connectivity. (**A-E**) Synaptic connectivity analysis exemplified in the rod pathway. Cross section (A) showing a ribbon synapse (open arrow) in a rod bipolar cell (RBC) axon terminal (green), a conventional synapse (filled arrow) in an A17 amacrine cell (AC) process (light blue) providing reciprocal feedback to the RBC, and an A2 AC process (in dark blue). 3D render of same synapse (B) and of involved cells (C). (D) Outputs (synaptic ribbons) and inputs (conventional synapses) of RBCs sorted by AC type (data from n=9 RBCs, each dot represents one cell pair; MF, medium-field). (E) Example skeletons of main AC types that synapse with RBCs (E), with Off and On starburst amacrine cell (SAC) stratification indicated (cyan and apricot, respectively). (**F**,**G**) Quantification of automatically detected ribbons in different bipolar cell (BC) types, with ribbon count (F, *left*), mean ribbon size (F, *right*), and distribution across IPL depth in (G, cyan and apricot lines indicate On and Off SAC plexuses, respectively; same BC dataset as in Fig. 4).

To scale this approach to the entire Eyewire II dataset, a deep learning model was trained to automatically detect synaptic ribbons using manually annotated ribbons as ground truth (for details, see Methods). We detected *≈*6.5 million ribbons in the volume and systematically evaluated ribbon statistics in BC types (Fig. 8F,G). Ribbon counts and average ribbon size of automatically detected ribbons both varied across BC types

(Welch ANOVA, *F* (14, 7818.37) = 15749.23, *p* < 0.001, *η*^2^*p* = 0.84; Fig. 8F). In general, RBCs had fewer but larger ribbons than cone bipolar cells (CBCs). Consistent with earlier work, Off CBC types t1, t2, and t7 had the highest ribbon counts (53). We found that CBC types t5t and t5o had significantly more ribbons than t5i (t5t vs. t5i, *p* < 0.001, *d* = 3.89; t5o vs. t5i, *p* < 0.001, *d* = 3.56; pairwise Games-Howell post-hoc test and Cohen’s *d* for independent samples with different variances; for full pairwise statistics see Fig. S3). Ribbon size varied more subtly, with Off CBCs typically having smaller ribbons and in particular the large CBC types t8, t9, and XBC having larger ribbons. Finally, ribbon density varied across IPL depth, with a somewhat higher density in the On vs. the Off sublamina, and decreasing density in the SAC bands. This analysis demonstrates the potential of Eyewire II, allowing large-scale analysis of retinal circuit motifs.

### Stranger Things in the retina

Besides synaptic circuits, the cellular ultrastructure resolved in Eyewire II allows us to study non-neural cells and other aspects of retinal cell biology in more detail. This includes, for instance, different types of glia, including contacts with blood vessels (Fig. 9A). Also, we reconstructed substantial parts of the vessel plexus in the volume (Fig. 9B). Since vasculature changes play a key role in retinal diseases like age-related macular degeneration and diabetic retinotopy (reviewed in 54), this makes Eyewire II well-suited for the joint investigation of retinal neurons, glia, and vasculature. The large volume offers also the opportunity to survey non-canonical types of neurons, such as cells with their soma situated within the IPL (Fig. 9C) or “chimeric” cells like ones that appear to possess feature of both ACs and BCs (Fig. 9D).

**Figure 9.**
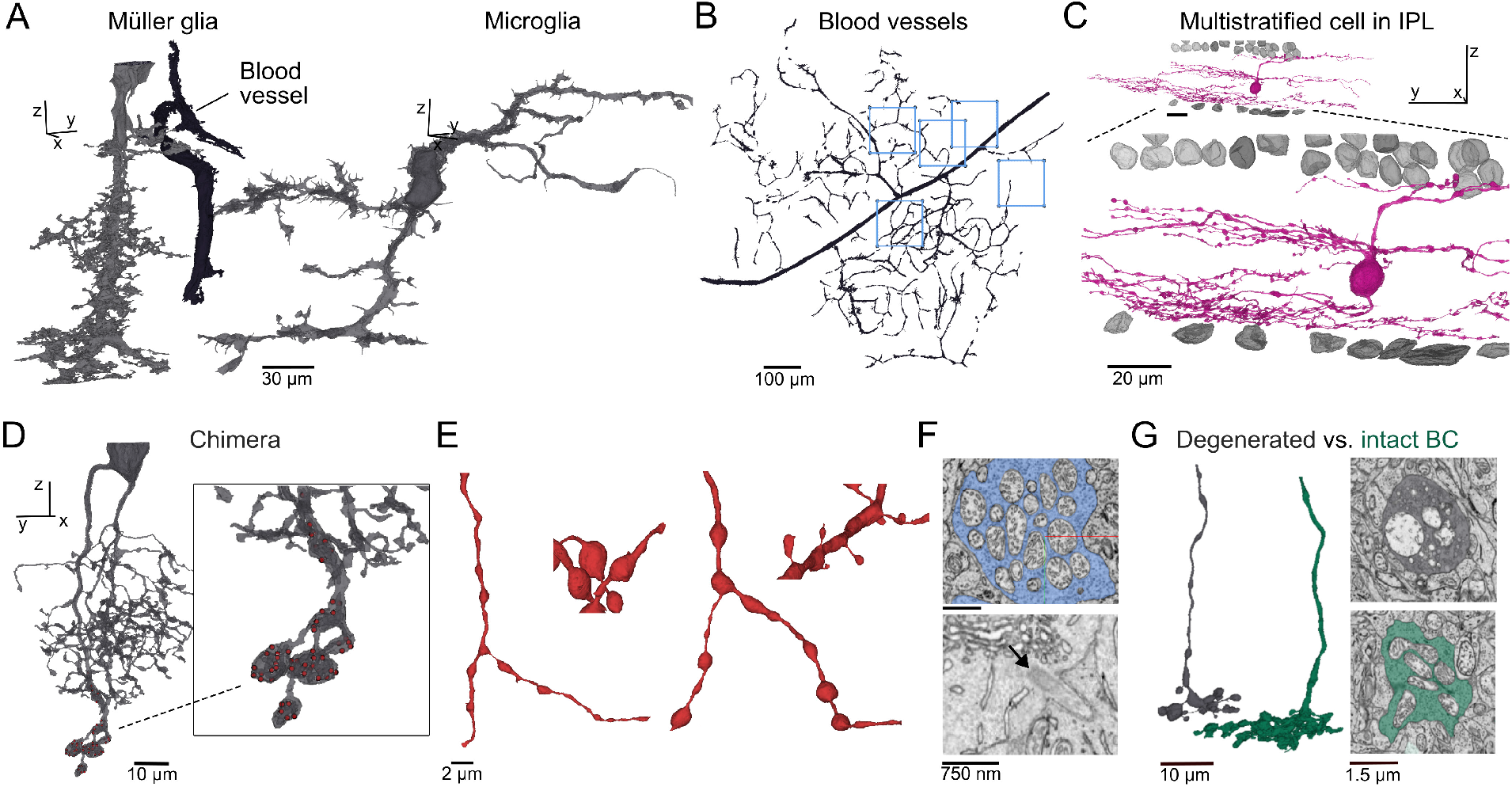
Other cellular structures. (**A**) Examples for non-neuronal cells in the EM volume. (**B**) Partial reconstruction of the vessel plexus (blue boxes indicate locations of recording fields). (**C**) Multi-stratified cell with soma within the inner plexiform layer (IPL). (**D**) Potential “chimeric” cell, with both small-field amacrine cell and bipolar cell (BC) properties (red dots indicate synaptic ribbons). (**E**) Examples for morphological specializations in retinal ganglion cell (RGC) dendrites (from *left* to *right*): Small-diameter bulbs, round bulbs, large-diameter bulbs, and thorn-like protrusions. (**F**) Exemplary intracellular structures: Mitochondria in an amacrine cell process (*top*) and primary cilium (arrow) in a RGC soma (*bottom*). (**G**) Example of degenerated t7 BC axon (gray, *top-left*) in comparison to a healthy counterpart (green, *bottom-right*).

In addition, sub-cellular morphological features can be studied: This includes the bulbous varicosities (“bulbs”) and/or small protrusions (“thorns”) present on the dendrites of many cells (Fig. 9E). Both bulbs and thorns varied in occurrence, shape, and size between cells, and could be implicated in synaptic processing. While they are well-known as features of starburst, A2, and A17 ACs, Eyewire II makes it possible to describe these features systematically across populations of retinal cell types. Many bulbs contained mitochondria (Fig. 9F, *top*), potentially enabling the study of metabolic demand in synaptic processing. At many RGC and AC somata, we observed primary cilia (Fig. 9F, *bottom*), which are little understood organelles implicated in neuromodulation (reviewed in 55). Finally, we also found examples of degenerating cells, which can be distinguished from their healthy counterparts by the structure of their mitochondria (Fig. 9G).

### The Eyewire II consortium

Achieving proofreading, celltype annotation, and synapse validation at the scale of Eyewire II is infeasible within a single laboratory. Large-scale connectomic reconstruction requires substantial person-hours for proofreading, but also expert domain knowledge to obtain ground truth annotations. Our team is supported by a growing consortium of over 30 labs from the field of retina research (eyewire.ai/consortium). Our community principles (eyewire.ai/principles) ensure fair credit for the work done by individual consortium members.

One of the most important achievements of this resource is the coordination of this community. Any lab can request to join the Eyewire II consortium through an online form (eyewire.ai/ew2_access). Once they agree to our community principles, our onboarding team trains them to view, proofread, and annotate the dataset, and provides material for self-guided practice. We host and distribute the data through a database of proofread and annotated cells that is regularly updated with the contributions of the community. For easy accessibility, we have developed a set of software tools (eyewire.ai/tools, Table 1), including an interactive cell browser and a command line tool for batch download of preprocessed cell morphologies. For both the database and the toolkit around it, we provide technical support through a community Slack workspace. Domain experts for curating ground truth are organized into specialized teams. Finally, we facilitate scientific collaborations by tracking ongoing projects and identifying shared research interests during regular town hall meetings where consortium labs present their preliminary work.

**Table 1.**
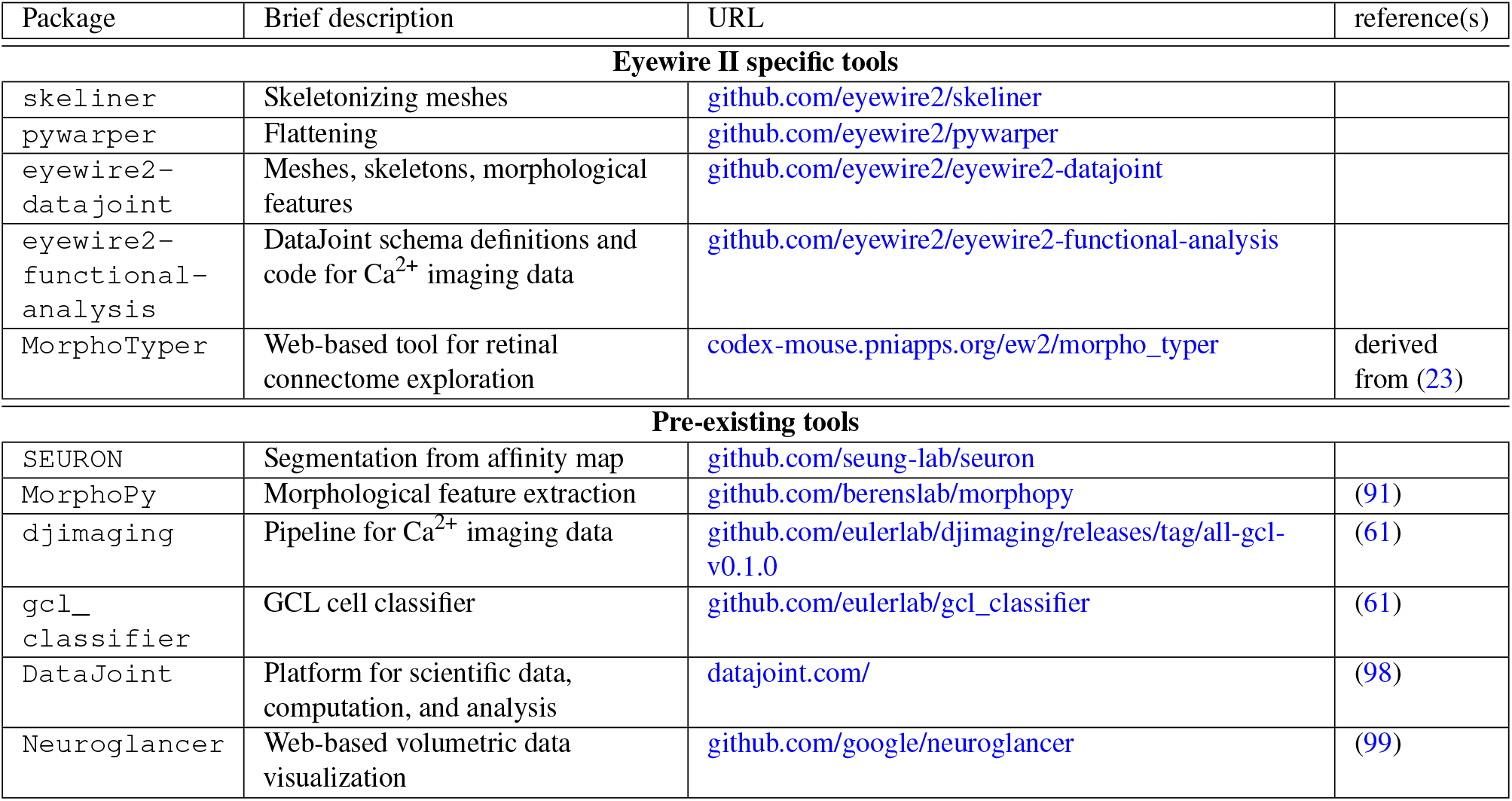
Tools. Overview of software packages employed for the curation and analysis of the Eyewire II dataset, including newly developed ones. All listed packages are open access.

## Discussion

Here we present Eyewire II, a large-scale EM dataset from the temporo-ventral mouse retina. It covers nearly 1 mm^2^ and is expected to contain *≈*100, 000 neurons, surpassing all prior retinal EM datasets by a factor of 10-100. With human proof-reading ongoing, we already reconstructed >25, 000 neurons, including wide-field AC types and complete mosaics of large RGC types. Unlike some earlier datasets (22, 25, 27), Eyewire II also includes synapses. An automatic synaptic ribbon detector allowed us to find almost all BCs (*≈* 42, 000). These cells were then classified into types based on axonal morphology, demonstrating the potential of Eyewire II for automatic morphological typing. Our human-in-the-loop cell-typing approach combined expert annotations with machine learning, and resolved all major retinal cell classes and numerous cell types, including candidate novel types. Furthermore, we showed that the rich set of light responses co-recorded with Eyewire II allowed us to link the morphological types in the GCL to well-known functional RGC types. Taken together, these results establish Eyewire II as a comprehensive resource for deciphering the cellular parts list and synaptic wiring of the vertebrate retina.

Key to understanding retinal circuits – like any other circuit in the central nervous system – is knowing the wiring diagram. Connectomics based on large-scale volumetric EM data is a very powerful approach (15–19), especially when combined with current machine-learning methods, web-based visualization tools, and an active online community (22). The power of this approach has been recently demonstrated for a cubic millimeter of mouse primary visual cortex (24) and the *Drosophila* brain (23).

Efforts to map retinal circuitry using EM began more than 50 years ago and have consistently provided crucial insights to the field (reviewed in 31, 56). At first, circuits that require only small volumes were targeted, laying the foundation of retinal connectomics by, for instance, mapping the midget pathway in the primate retina (e.g., 57) and uncovering the anatomical basis for interactions between rod and cone signaling (e.g., 13). Later, techniques such as block-face scanning EM (15) provided larger volumes, which opened up the possibility to assess circuit motifs systematically. This enabled, for example, revealing the synaptic wiring underlying direction selectivity (21, 25, 58) and extracting structural motifs, such as the pathways through which ACs suppress rod signals in cone BCs (10) and the synapse configurations present at the axon terminals of different BC types (53).

Despite these efforts, our understanding of most retinal circuits remains incomplete. One critical limitation has been the restricted size of available EM datasets: The largest datasets, such as RC1 in rabbit (28), and e2006 (25) and e2198 (21) in mouse, cover around 0.05 to 0.1 *mm*^2^ (see Table 1 in 29), which hampers the reconstruction of rare cell types and cells with large arbors (cf. Fig. 1A). Another major limitation has been the time effort needed for curation – segmentation, proofreading, and annotation of the cells. For instance, proofreading the 400 RGCs in Eyewire I (22) took a community of thousands of volunteers about a year, with approx. 9, 250 primary proofreading actions required per RGC. Despite differences in proof reading procedures between Eyewire II and Eyewire I, the average RGC in Eyewire II required only 55 proofreading edits, thanks to advances in machine-learning algorithms used for segmentation. This reduced the primary proofreading effort by a factor of*≈*170, and is a critical ingredient to enable large-scale reconstruction.

A central requirement for any connectomics resource is to enable researchers to rapidly identify cells of interest. Achieving this goal at the scale of Eyewire II– with tens of thousands of reconstructed neurons – demands an automatic approach building on available domain expertise. We addressed this challenge through a human-in-the-loop strategy that iteratively combined expert annotation with supervised machine learning classifiers and unsupervised clustering. We demonstrate the full potential of this approach for BCs, where the combination of axonal IPL depth and morphology provided a sufficiently rich feature space to classify all 15 known BC types (27, 36– 38) with near-complete mosaics. Distinguishing BC types t8 and t9 was a challenge for the classifier, as these types are very similar with respect to terminal morphology and IPL stratification depth. Here, adding dendritic arbor features is expected to improve separation, because t9 selectively contacts “true” S-cones, whereas t8 contacts both cone types (38, 59). Currently, the Eyewire II dataset contains BC dendritic arbors only in a small patch (see t5i, t8, and t9 examples in Fig. 4C); this coverage will increase as imaging proceeds.

For RGCs, the almost complete reconstruction of all cells in the GCL provides a strong basis for future automatic classification. Complete classification of ACs may present a greater challenge, as super-wide-field polyaxonal types (e.g., 39) with sparse arbors and other large ACs reach the volume boundaries. However, most AC types are small and many examples of these types are completely contained within the volume. For example, for the SACs almost a thousand examples are contained in the volume. Therefore, we expect that Eyewire II will critically advance AC classification. For both RGCs and ACs, mosaic regularity will serve as a powerful validation criterion (60). In addition, internal morphological features – such as mitochondrial organization and the fine morphological specializations – as well as connectivity profiles are expected to provide additional sources of validation. Taken together, the Eyewire II dataset offers a scalable path toward a comprehensive, community-curated cell atlas of the mouse retina.

Another strength of Eyewire II is that morphological reconstruction is paired with light responses for a subset GCL neurons. This makes it possible to examine how morphological cell type maps onto functional response type, and where the two classifications converge or diverge. Using the morphological and functional data as anchors, allows integration with further datasets and modalities: For example, a recently published mouse GCL dataset (61) provides responses of *≈*80, 000 GCL cells to visual stimuli, which can be used for a fine-grained functional analysis. In addition, matching dye-filled cells from patch-seq recordings of RGCs (46) to their Eyewire II counterparts, will allow integration with transcriptomic resources (5). A similar strategy could be used to achieve an integrated multimodal type classification for ACs, where morphological, transcriptomic (2, 6, 62) and functional diversity (63, 64) is known to be extensive.

A prerequisite for mapping retinal circuits is the reliable, large-scale identification of synapses throughout the reconstructed volume. Unlike several earlier mouse retinal EM datasets, which intentionally omitted intracellular staining (e.g., 25), Eyewire II was processed to preserve intracellular ultrastructure, rendering synaptic ribbons and conventional synapse vesicle clouds clearly visible. A first automatic machine-learning detector for ribbons found nearly 6.7 million ribbons, which allowed us to systematically compare ribbon counts and size across BC types. We confirmed, for instance, that RBCs have fewer but larger ribbons than CBCs (65), consistent with the distinct release dynamics and temporal filtering properties of these two pathways (66) and with differences in ribbon architecture and temporal dynamics in rods and cones (67, 68). Intriguingly, we also found that CBC types t5t and t5o have significantly more ribbons than t5i. This difference may reflect distinct presynaptic gain settings (69). Overall, counts of our automatically detected ribbons match manual counts from previous studies for most BC types, but are slightly higher for t1 BCs (53, 70, 71). Extending these approaches to automatically detect not only ribbon-associated synaptic sites but also conventional synapses will enable comprehensive quantification of both excitatory and inhibitory connectivity across the entire IPL. Reconstructing such a large-scale connectome of the inner retina will pave the way towards systematic identification of cell-type specific connectivity rules and circuit motifs underlying retinal computations. This fine-grained information about each cell’s connectivity will further improve automatic cell typing, and enable computational models of the retina that make experimentally testable predictions about the functional significance of circuits and cell types (72, 73).

There are a few limitations of the current dataset that merit discussion. First, the EM imaging does not yet extend fully to the OPL, meaning that outer retinal circuits cannot be comprehensively studied in Eyewire II at present, and that BC dendritic arbors and HCs are only available in a small sub-region. Imaging is ongoing and is expected to eventually provide OPL coverage in parts of the volume, enabling reconstruction of the full vertical circuit from photoreceptors to RGC outputs.

Eyewire II will then also allow studying mosaics of complete HCs, including their extended, offset axon terminal system (74) systematically at the ultrastructural level (e.g., 75). Second, functional Ca^2+^ data are available from only *≈*400 GCL cells. This implies limited coverage for large types, of which only a few examples were recorded, limiting how broadly morphology-function relationships can be generalized. Related to this, the sequential recording of neighboring fields led to progressive light adaptation across fields, a confound that will need to be considered in functional analyses. Third, gap junctions, which mediate electrical coupling between many retinal cell types (76), are not visible in the Eyewire II volume. Given that the EM protocol was not optimized to preserve extracellular space and the spatial resolution of EM imaging, inferring candidate gap junction sites is difficult (77). To overcome this, a potential starting point is the identification of apposition points between cell type pairs known to be coupled (e.g., A2 ACs and CBCs; 69, 78), followed by testing for the absence of synaptic markers to generate ground truth data for network-wide analysis.

The analyses presented here are intended as a proof of principle for what the Eyewire II dataset will enable as proofreading and annotation continue. The most immediate extensions will be to complete cell typing of RGCs and ACs, and automate detection of post-synaptic ribbon partners and conventional synapses. Together, these advances will bring a comprehensive wiring diagram of the mammalian retina within reach. Beyond connectivity, the preservation of intracellular ultrastructure in Eyewire II opens the door to volumetric analysis of organelle organization, including mitochondrial distribution and the relationship between organelles and synaptic active zones. These and other subcellular features may vary systematically across cell types or reflect metabolic state. The dataset also provides the foundation for a complete cell atlas of the mouse retina, integrating morphological, functional, and genetic identity across all neuronal classes. Finally, the open, community-based model of the project – combining professional proofreaders, domain experts, and citizen scientists alongside web-based visualization and analysis tools – offers a template for how datasets of this scale can be made broadly accessible and continuously improved by the wider research community.

## Methods

### Experimental procedures

#### Animal procedures

One male, approx. 3-month-old wild-type mouse (C57Bl/6J, JAX 000664), purchased from Jackson Laboratory, was used in this study. Before the experiment, it was housed under a standard 12 h day/night cycle at 22°C and 55% humidity. All animal procedures were approved by the governmental review board (Regierungspräsidium Tübingen, Baden-Württemberg, Konrad-Adenauer Strasse 20, 72072 Tübingen, Germany) and performed according to the laws governing animal experimentation issued by the German Government.

#### Tissue preparation for Ca2+ imaging

Before preparing the retina for EM, visual responses of a subset of GCL neurons were recorded with 2p Ca^2+^ imaging – as established in earlier work (34). The mouse was dark-adapted for *≈*1h before tissue preparation; all following experimental procedures were carried out under very dim red light (> 650 nm). Next, the animal was anesthetized with isoflurane (Baxter) and killed with cervical dislocation. Next, the eyes’ orientation was marked before they were quickly enucleated in carboxygenated (95% O_2_, 5% CO_2_) artificial cerebral spinal fluid (ACSF) solution containing (in mM): 125 NaCl, 2.5 KCl, 2 CaCl_2_, 1 MgCl_2_, 1.25 NaH_2_PO_4_, 26 NaHCO_3_, 20 glucose, and 0.5 L-glutamine (pH 7.4). After removing cornea, sclera, and vitreous body, the retina was flattened on an Anodisc (#13, 0.2 *µ*m pore size, GE Healthcare) with the GCL facing up. We then loaded the retinal tissue with the fluorescent Ca^2+^ indicator dye Oregon-Green BAPTA-1 (OGB-1, hexapotassium salt, Life Technologies) using electroporation (79). To this end, we placed the Anodisc with the tissue and a drop of 5 mM OGB-1 in ACSF between a pair of platinum electrodes (CUY700P4E/L, Nepagene/Xceltis) and applied voltage pulses (11 pulses, +9.5 V, 100 ms pulse width, at 1 Hz) from a pulse generator/wide-band amplifier combination (TGP110 and WA301, Thurlby handar/Farnell). Finally, the Anodisc with the tissue was transferred to the recording chamber of the microscope, where it was continuously perfused with carboxygenated ACSF (at 36°C and 4 ml/min) and left to recover for *≈*60 min before the recordings started. To visualize the tissue, the ACSF contained *≈*0.1 *µ*M Sulforhodamine-101 (SR101, Invitrogen).

#### Two-photon Ca2+ imaging

We used a MOM-type 2p microscope (Sutter Instruments/Science Products) as described previously (80, 81). Its excitation source was a mode-locked Ti:Sapphire laser tuned to 927 nm (Mai Tai-HP DeepSee, Newport Spectra-Physics) and it featured two fluorescence detection channels, “green” (HQ 510/84, AHF) and “red” (HQ 630/60, AHF) to visualize OGB-1 and SR101, respectively, through a water immersion objective (W PlanApochromat 20×/1,0 DIC M27, Zeiss). For image acquisition, we used a custom-made software (ScanM, by M. Müller, MPI, Martinsried, and TE) running under IGOR Pro 6.3 for Windows (Wavemetrics), with 64× 64 pixel frames at 7.8125 Hz for time-elapsed recordings and 512 × 512 pixel frames for higher resolution images. In both configurations, recording field size was 94 × 94 *µ*m.

We recorded from five fields in the temporo-ventral retina (F0 to F4; Fig. 6A-C). The retinal position of each field, as well as the positions of optic disc and retinal edges were logged, which enabled locating the fields in the embedded tissue (Fig. 6C). The high-resolution images allowed us to register the recorded GCL cell somata with the segmented ones (e.g., Fig. 6C vs. D) – aligning morphology (Fig. 6E,F) and Ca^2+^ responses (Fig. 6G).

#### Light stimulation

Light stimuli were projected through the objective lens (80) and centered to the recording field using a digital light processing (DLP) projector (LightCrafter E4500 MKII, EKB Technologies Ltd.) supplied with light from external, band pass-filtered LEDs through a common light-guide (green: 576 BP 10, F37-576; UV: 387 BP 11, F39-387; both AHF/Chroma; for details, see 82). This allowed us to differentially excite the two spectral mouse cone types (M/PR1 and S/PR4) with their peak sensitivities at 510 and 360 nm (83), respectively. Stimulator intensity (as photoisomerization rate, 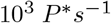 per cone) was calibrated to range from *≈*0.5 to *≈*20 for M-(LWS) and S-(SWS1) opsins, respectively. We estimate that a steady illumination of 1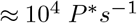 per cone was present during the recordings due to the 2p excitation of photopigments by the scanning laser (81).

Visual stimuli were generated and presented with QDSpy (github.com/eulerlab/QDSpy). To capture different aspects of visual processing, we presented three stimuli, which probe temporal, motion-, and naturalistic pattern-related response properties:

1. An achromatic full-field stimulus consisting of a bright step and two sinusoidal intensity modulations, one with increasing frequency (0.5 to 8 Hz) and one with increasing contrast (“full-field chirp”; 34).
2. An achromatic bright-on-dark bar (1, 000 *µ*m long, 300 *µ*m wide) moving at 1 mm/s in eight directions (“moving bar”; 34).
3. A dichromatic natural movie (Fig. 6G right) that consisted of 113 5s clips from taken from footage recorded outside in the field using a custom camera (35) that captured the spectral bands relevant for mouse vision. Five of the clips represented the “test set” and were repeated trice – at the beginning, the middle, and the end of the stimulus – to enable assessing the response reliability of the neurons. The remaining clips were displayed in pseudo-random order. The stimulus design followed that of Höfling et al. (84), with modifications: Clip transitions were not explicitly balanced to control for brightness changes, and the set of clips was partially altered.

#### Tissue fixation and EM staining

After the Ca^2+^ imaging experiment, the tissue was immediately fixed overnight in 4°C phosphate buffer containing 2% PFA (paraformaldehyde, EMS Diasum) and 2% GA (glutaraldehyde, EMS Diasum). The following morning, the tissue was rinsed 4 times for 30 minutes in 4°C phosphate buffer and subsequently stained and fixed in 2% osmium tetroxide (OsO_4_, EMS Diasum), reduced with potassium ferro-cyanide (Sigma Aldrich) and buffered in 0.1 M cacodylate (EMS Diasum) buffer for 2 hours on ice in the dark. Prior to staining, the reduced osmium solution was filtered with a 0.2 *µ*m syringe filter (Fisherbrand). After the reduced osmium step, the tissue was washed 4 times for 15 minutes in ultrapure water on a rotor at room temperature except for the last washing step, where the sample was placed in an oven at 60°C. At the same time, the thiocarbohydrazide (TCH, EMS Diasum) solution was prepared in the 60°C oven.

Next, the sample underwent a concentration gradient: every 4 minutes, the TCH concentration was increased starting at 0.25%, then 0.38%, 0.72%, and ending at 1% TCH. The double distilled water for the first post-TCH washing step was also heated to 60°C. After the first wash in heated water, the temperature was transitioned to room temperature again and subsequent washing steps (to reach 5 times 12 minutes) were continued at room temperature. The tissue was further post-fixed and stained in a 2% OsO_4_ solution for 45 minutes. To increase unspecific protein labeling, the tissue was additionally stained with uranyl acetate and lead aspartate. Prior to the uranyl acetate staining step, the tissue was washed in maleate buffer (0.05 M, at pH 5.5 with NaOH, both EMS Diasum) 5 times for 24 minutes and then stained in 1% uranyl acetate in pH 6.0 maleate buffer overnight.

The next day, the sample was washed 5 times for 24 minutes in double distilled water and transferred into a 60°C lead aspartate solution for 30 minutes (0.132 g lead aspartate in 20 ml 0.03 M aspartic acid solution at pH 5.3; lead nitrate from EMS Diasum, aspartic acid from Sigma-Aldrich). The tissue underwent four 30-minute washes in ultrapure water. Dehydration then proceeded through a graded ethanol series (20%, 50%, 70%, 100%, 100%, 100%; Sigma-Aldrich), followed by 100% propylene oxide (EMS Diasum), with each step lasting 10 minutes. Finally, Epon resin (Epon 812, Ladd Research and EMS Diasum) was slowly mixed into the propylene oxide (2 : 1, 1 : 1, and 1 : 2, each for 1 hour) and the samples were then transferred into 100% Epon, first for a half an hour, then for 1 hour with freshly made Epon and another two rounds with the same batch of fresh Epon. Once those infiltration steps had concluded, the sample was embedded and cured in the oven for a week at 60°C.

#### Sectioning, imaging, and alignment

Ultra thin sections were cut at 40 to 45 nm on a UC7 ultramicrotome (Leica). Sections were collected onto carbon nano tube tape (85) with the help of the ATUM (86). Subsequently, a modified version of the WaferMapper software (86) was used to drive imaging of the sections on a Merlin double condenser scanning electron microscope (Zeiss) with the help of an external scan generator and acquisition software (smartSEM 6.04-7.05 and API, Zeiss; Atlas and API, Fibics). 17× 17 tiles with 6% overlap were acquired at 800 ns dwell time with a field of view of 65.536 *µ*m and 8 nm resolution. Modification of the WaferMapper software specifically included sub-degree alignment of the sections before imaging to reduce the number of tiles that had to be acquired.

Once the tiles were acquired, stitching was done with MATLAB (MathWorks) and uploaded with the CloudVolume package (87). Fine alignment of the dataset has been performed by Zetta AI (zetta.ai/).

### Data analysis – Anatomy

#### Automated segmentation

Segmentation (Zetta AI) was performed as described by Kim et al. (88) on a down-sampled 16× 16 nm version of the dataset. In brief, a network was trained to predict three distinct output heads: (1) 24-dimensional dense voxel embeddings, (2) nearest neighbor intervoxel affinities, and (3) a myelin mask, with the data foundation of the model being publicly available ground truth data (Microns consortium; 24). For fine-tuning, manual ground truth segmentation was used in small volumes from (88) and for this, mitochondria were deliberately over-segmented to reduce merge errors, accepting an increased risk of split errors as a trade-off. For segmentation, affinities within the myelin mask were masked out and not used for subsequent segmentation steps. Other than in (88), we applied a glia detection for the Müller cell end feet. We then used the distributed agglomeration algorithm described in (89) to create the segmentation from the affinity map. The inference and segmentation was done by SEURON (see Table 1)

#### Human proofreading

Eyewire II uses CAVE for hosting the proofreadable segmentation and annotations (90). Proofreading procedures and workflow were similar to those previously described (23). Human proofreaders consisted of a group of 28 researchers, 72 paid and trained specialists, and 11 volunteer citizen scientists. On-boarded researchers underwent proofreading training and completed self-guided tutorials and a final assessment before obtaining edit-access for proofreading. Paid professionals and citizen scientists had prior experience from proofreading other EM datasets. To obtain quality control cells were regularly proofread by multiple paid professionals, and techniques such as collaboration on difficult problem areas and deference to promoted experts helped ensure a robust dataset.

#### Ribbon synapse detection

Automated ribbon localization (Zetta AI) was based on manually annotated ribbons from four sub-volumes used as ground truth. These included one large volume (14× 20 *µ*m) vertically spanning the full IPL and four small volumes (8× 8 × 3, 88 × 8, and twice 16× 16× 8 *µ*m), which were randomly scattered across the dataset.Ribbons were manually annotated in every section in which they were visible. After initial training, model predictions were reviewed by expert annotators; missed true ribbons were incorporated into the ground truth and false detections were labeled accordingly. Then, the model was retrained on the updated dataset.

The final annotated ground-truth contained *n* = 5, 785 annotations. Final performance was assessed on the held-out test set with *n* = 384 annotations, which was refined after the initial review but remained excluded from all training steps. For downstream analysis, automatically detected ribbons smaller than 100 voxels or located outside the segmentation mask were filtered out.

#### Ribbon statistical analysis

Significant differences in ribbon counts and sizes across types were assessed using Welch’s ANOVA, which accounts for unequal variances across groups. Pairwise statistical differences of ribbon count and size was determined using post-hoc pairwise Games-Howell tests using the Python `pingouin` toolbox to calculate *p*-values and Cohen’s *d* for effect sizes. Ribbons with a z-depth above 25 in SAC-band normalized space where excluded to remove dendritic ribbons.

### Data analysis – Morphological cell typing

#### Skeletonization and flattening

Center-line skeletons were extracted from the reconstructed 3D meshes using `skeliner`. Briefly, each mesh is partitioned into thin geodesic shells from the soma, each shell is clustered into one or more interior nodes carrying a local radius estimate, and mesh edges are projected onto the resulting node graph; soma-like nodes are then collapsed, disconnected patches are bridged, and a minimum-spanning tree is computed to yield an acyclic skeleton rooted at the soma. The resulting skeletons were then flattened into a common inner-plexiform-layer (IPL) coordinate frame using `pywarper`, a Python re-implementation of the conformal-mapping pipeline of Sümbül et al. (40). The ON and OFF SAC surfaces were estimated from the dendritic skeleton nodes of reconstructed SACs: for each layer, *z* was sampled on a common 5 *µ*m XY lattice as the median depth of all nodes within one grid spacing of each lattice point, and remaining gaps in the OFF layer were filled by inverse-distance-weighted interpolation of the local ON–OFF offset. The two layers were then fit to smooth height fields and flattened by a quasi-conformal map that straightens the SAC surfaces while minimizing local angular distortion. Each skeleton node was re-registered into this frame by a local polynomial least-squares fit referencing both SAC layers, preserving its relative depth between the bands. In the resulting warped skeleton, the *xy*-plane is parallel to the SAC layers and *z* corresponds to IPL depth, with *z* = 0 and *z* = 12 defined as the ON and OFF SAC bands.

#### Density profiles

From each warped skeleton, we computed three-dimensional neuritic density profiles in cylindrical coordinates (*r, θ, z*) centered on the soma; for RGCs we excluded the axon. Each edge of the skeleton was assigned a mass equal to its length, and these masses were accumulated into a 3D histogram on a user-defined grid. Each bin was then divided by its physical volume 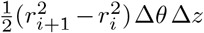 to yield a dendritic length density (µm/µm^3^). To separate structure at different orders of spatial dependence, we normalized each density to a probability distribution *p*(*r, θ, z*) and decomposed it into 1D marginals *p*(*r*), *p*(*θ*), *p*(*z*) and 2D marginals *p*(*r, θ*), *p*(*r, z*), *p*(*θ, z*). Pairwise residuals were defined as the deviation of each 2D marginal from the product of the corresponding 1D marginals (i.e., the distribution expected under independence), e.g. *p*(*r, θ*) *−p*(*r*) *p*(*θ*), isolating the component of joint structure not explained by the two axes independently.

#### Features

Morphological features were computed using custom Python functions and `MorphoPy` (91). Features included: (*i*) number of tips and branch-points, (*iii*) principal components from weighted functional PCA of the 1D marginals *p*(*r*), *p*(*θ*), *p*(*z*) and the 2D residual profiles, (*iv*) mean of the (unnormalized) 3D density *d*(*r, θ, z*), (*v*) percentiles of the 1D z density profile (for visualization only), (*vi*) z-position of the soma, (*vii*) median tortuosity, (*viii*) summary statistics of branch angles and path angles (between neighboring nodes without a branch point), (*ix*) summary statistics of neurite radii, and (*x*) median path length of intermediate and terminal segments.

#### Expert annotations

Cells type and class assignments were updated iteratively during the curation of this dataset (see also Human-in-the-loop). For this we relied on expert annotations that were guided using machine learning tools including supervised classification and unsupervised clustering. Cell information was organized in a shared spreadsheet, where different experts contributed to label cell class and type information to proofread cells. To facilitate this process, we developed `MorphoTyper` (see Table 1), a web-based tool specifically for retinal connectome data exploration, derived from FlyWire Codex (23).

#### Classification

To determine the class and type of proofread cells, we used random forest classifiers from the Python package *scikit −learn* (92). Human expert labels were split into a training, calibration, and test set, using stratified sampling, with special handling for singleton classes, which were retained in the training set. The random forest classifiers were trained with 500 estimators on the training set using balanced class weights to account for imbalanced class distributions. To obtain well-calibrated probability estimates, we applied sigmoid calibration using `CalibratedClassifierCV` (from the `sklearn.calibration` package) on the calibration set. The classifier labels and the respective confidences were then provided back to the human expert annotators. As the dataset and the number of available annotations grew, we iteratively updated the classifiers.

#### Clustering

Graph-based clustering was performed using the Leiden algorithm (93) implemented in scanpy (94) to obtain both RGC and BC clusters that we compared to human annotations. We used these clusters to detect outliers, propose new types and to propose candidate cells for known cell-types. To this end, we implemented a pipeline that leverages both cluster membership and nearest-neighbor connectivity.

For each known cell type, we identified outlier cells belonging to singleton clusters or clusters where that type represented *<* 15% of the population. We then identified “majority clusters”, which contained either at least 3 cells of the target type or where that target type comprised > 40% of the cluster. Unlabeled cells within majority clusters were proposed as high-confidence candidates, and mislabeled cells were proposed for relabeling when outnumbered by the correct type. For cell types with fewer than 5 candidates, we performed connectivity-based search using the nearest-neighbor graph, identifying unlabeled cells with edge weights > 0.3 to non-outlier reference cells. When sufficient reference cells existed, we required cumulative connectivity weights of≥ 1.0. Candidates were cross-validated against other cell types and excluded if they showed 1.5 times stronger connectivity to an alternative type. Finally, clusters not assigned to any known cell type and containing at least 10 cells with *<* 50% labeled were flagged as potential novel cell types.

#### Bipolar cell typing

To identify most of the BCs in the volume, we collected all segments (*n* = 46, 981) with a ribbon count > 35 in the IPL (defined as *z* < 25); this threshold is based on ribbon counts in manually identified BCs. Through manual inspection of subsets, we confirmed that most of these segments were indeed BCs: Since the majority of these had not been proofread yet, we applied a combination of quality filters to remove cells that were incomplete or still required significant proofreading. Specifically, we removed cells with a height (*max*(*z*) *− min*(*z*)) *<* 12 *µ*m to reject cells too close to the upper boundary of the volume. Further, we removed cells that had incorrect neurites attached, by computing a convex hull for each cell, and filtering all cells with a perimeter *p >* 300, a diameter *d >* 80, or *p/*(*dπ*) > 1.35.

We then clustered the remaining cells (*n* =42, 000) using Leiden clustering (neighbors= 5, resolution= 2). In each cluster, we removed cells that were outliers in hull perimeter using a robust z-score approach. Specifically, for each cluster we computed the median hull perimeter 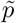 and the inter-quartile range *IQR*, and scored each cell by its deviation from the cluster median, both in raw units 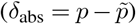 and normalized by the 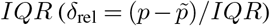. Cells (*n ≈* 300) were excluded if |*δ*_rel_| > 2.5 or |*δ*_abs_| > 30.

Similar to previous studies (27, 38, 95), we used stratification profiles as the main feature together with a few other morphological features. As most of the BCs were incomplete (i.e. lacking dendrites and, in many cases, soma) due to the boundaries of the reconstructed volume, we decided clip all BC skeletons above *z* = 25 to align them better. Features were normalized and clustered using the Leiden algorithm at two resolutions: a coarse partition (*k* = 20 neighbors, resolution= 1) and a fine partition (*k* = 10 neighbors, resolution= 5), resulting in 18 and 80 clusters, respectively.

To identify coarse clusters containing multiple types we computed the coverage for all coarse clusters. Coarse clusters exceeding a mean count of 1.5 cells in occupied areas were flagged and re-clustered using the fine Leiden partition, replacing their coarse cluster label with the corresponding fine-grained assignment. The resulting set of clusters was then subjected to a iterative merging procedure, in which pairs of clusters were merged if they were sufficiently similar in hull diameter, dendritic depth (characterized by the 5^*th*^ depth percentile) and the normalized feature space. This resulted in 15 merged clusters, each corresponding to one of the known BC types. After that we did one final step of human expert refinement (also making use of the mosaics), to remove e.g. obvious mosaic violating cells and to better separate t8 and t9 BCs. As described above, we then trained a random forest classifier to predict type labels for both labeled and unlabeled BCs.

#### Consensus labels

The cell class and type labels we report here are a combination of expert labels and classifier predictions. For each cell, we derived a consensus label for both cell class and cell type by reconciling the human annotation with the classifier output using a four-way decision scheme. Each consensus assignment was tagged with a decision category indicating how the label was determined: *Both strong* (human and classifier agree, with the classifier highly confident), *both weak* (human and classifier agree, but the classifier is less confident), *classifier* (no human label available, but the classifier is highly confident), or *conflict* (human label present but contradicted by the classifier).

For cell class, we considered the three classes AC, RGC, and BC, comparing the human label against the calibrated classifier probability for that class. A cell was assigned a class as *both strong* if the human label matched and the classifier probability for that class was > 0.75, and as *both weak* if the human label matched and the classifier probability fell between 0.25 and 0.75. Cells without a human class label were assigned the classifier’s prediction (decision category *classifier*) whenever its probability exceeded 0.75. Cells whose human label was contradicted by a classifier probability *<* 0.25 were flagged as *conflict* and received no consensus class. As BCs are defined by their ribbon-type output synapses in the IPL, we additionally required any cell assigned to the BC consensus class to have > 35 synaptic ribbons within the IPL; cells failing this criterion were stripped of their BC consensus assignment and tagged *no ribbons*.

For cell type, we compared the human-annotated type against the classifier’s three most probable predictions and their associated probabilities. A cell was labeled *both strong* if the human type matched the classifier’s top prediction and that prediction carried a probability > 0.75. It was labeled *both weak* if the human type appeared among the top three predictions with a probability between ≥0.1 and 0.75. Cells without a human type label were assigned the classifier’s top prediction (decision category *classifier*) whenever that prediction’s probability exceeded 0.75. Cells whose human label did not appear among the top three predictions at probability ≥ 0.1 were flagged as *conflict* and received no consensus type.

Only cells assigned a decision category of *both strong, both weak*, or *classifier* were considered to carry a valid consensus label for downstream analyses.

### Data analysis – Ca^2+^ responses

#### Regions of interest

Regions of interest (ROIs), which correspond to the somata of individual cells in the GCL, were manually drawn in the input data using IGOR Pro. Next, raw Ca^2+^ traces were extracted from individual ROIs as the mean over all ROI pixels for each imaged frame.

#### Signal processing

Raw traces were detrended by subtracting a smoothed estimate of the signal (*r*_*smooth*_), removing slow, non-stimulus-related drifts while preserving the light-evoked fluctuations for further analysis. The smoothed trace *r*_*smooth*_ was computed by applying a Savitzky-Golay filter (96) of 3^*rd*^ polynomial order and a window length of 60 s using the Python `SciPy` implementation (97).

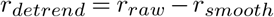

For the chirp and moving bar stimulus, we computed response averages from the detrended traces as means over repetitions.

In the case of the moving bar stimulus, we also computed a response average projected to the preferred motion direction of the cell (for details, see 34). Finally, response averages were normalized by first subtracting the baseline activity (computed as the mean over the first second), and then by dividing by the maximum amplitude *max*_*t*_(|*r*(*t*)|) = 1. This normalization was performed independently for each ROI and stimulus.

#### Response quality

To quantify the response quality of individual ROIs, we computed a quality index (*QI*) for chirp (*QI*_*chirp*_) and moving bar (*QI*_*MB*_) stimuli:

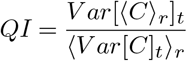

where *C* is the *T* by *R* response matrix (time samples by stimulus repetitions) and ⟨ ⟩_*x*_ and *V ar* []_*x*_ denote the mean and variance across the indicated dimension *x*, respectively. For the moving bar, *QI* was computed for all directions independently using the direction with the largest *QI* as *QI*_*MB*_. Cells with either *QI*_*MB*_ > 0.6 or *QI*_*chirp*_ > 0.35 were considered *responsive* in the following analysis.

#### Functional classification

We predicted the functional type of all cells from their mean Ca^2+^ responses using a previously published GCL type classifier (61), trained to predict the functional types described earlier (34). The classifier outputs soft labels, assigning each cell a probability of belonging to each functional type. We incorporated morphology-based cell-class constraints by setting the probability of functional AC types to zero for RGCs (and vice versa) and re-normalizing the resulting distributions.

#### Estimating stimulus exposure

To estimate how strongly the cells in a recording field were *activated* by the stimulus history, we calculated activation *a*(*t*) as as a leaky integrator:

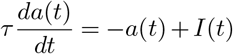

with time constant *τ*, taking into account that the cells adapt to the stimuli (and “recover” in breaks between stimuli), and stimulus intensity *I*(*t*). For simplicity, we only considered the stimulus history from the first stimulus presented to F0 to right after the presentation of the last stimulus to F4. We did not include the exposure by the excitation laser, as this was roughly similar for each field.

## Code & data availability

All code repositories developed for this study are collected in the GitHub organization eyewire2 (github.com/eyewire2), including a repository with code to reproduce all figures and analysis presented in this work. Some packages are already public, all other packages relevant to this resource paper will be made public at the latest upon journal publication (for an overview, see Table 1).

For access to the dataset, anyone can request access via our sign-up website (eyewire.ai/ew2_access) and will be offered an onboarding as soon as possible. This includes full access to EM images, segmentation, meshes, and skeletons through the web-based interface Neuroglancer, as well as access to the Ca^2+^ imaging data (for instructions, see GitHub repository github.com/eyewire2). In addition, the data presented and analyzed in this manuscript will be made available on Huggingface (huggingface.co/datasets/eulerlab/eyewire2-data), at the latest by the time of journal publication.

### EM data

To process the meshes, several Python packages were developed, which are all available on GitHub (for details, see Table 1): `skeliner` was used for skeletonizing meshes, which were then flattened using `pywarper`. Meshes, skeletons, and derived morphological features were organized in a custom DataJoint (98) database. A subset of morphological features was computed using the previously published Python package `MorphoPy` (91).

### Ca2+ imaging data

To process the Ca^2+^ imaging data, several Python packages were developed, which are all available on GitHub (for details, see Table 1): The Ca^2+^ imaging data was processed as described earlier (61). Briefly, the data was organized in a custom DataJoint (98) database derived from the Python package `djimaging`. The exact schema definition, i.e. which `djimaging` tables have been used and how, and all code to reproduce the Ca^2+^ related figures of this manuscript, is available from the `eyewire2` repository. The functional classification of GCL cells was done using `gcl_ classifier` from Gonschorek et al. (61).

## ACKNOWLEDGEMENTS

We dedicate this work to the memory of Richard H. Masland, whose insight and passion for the retina were an enduring source of inspiration. We thank Adrian A Wanner, Eric W Hammerschmith, Christel Genoud, and Ashwin Vishwanathan for helpful discussions and Katherine Nelson† and Sylvia Bolz for excellent technical assistance. We thank Christian Puller for helpful advice on ribbon identification. We thank Eyewire II community members Seunghoon Lee, Yao Xue, Dario Tommasini, Kali Mathura, Selma Tahri, Kacey Azizi-Namini, Ritsuko Fujimoto, and all the Eyewire II Citizen Scientists – in particular Anne Kristiansen, Andrea Becker, Tom Stocks, Krzysztof Kruk, Jaime Skelton, Matthew Lichtenberger, *Mavil*, Ashley Morren, – for their proofreading and annotation efforts. Eyewire II was funded by Princeton University (PNI; to HSS), the National Institutes of Health (NIH; U01 NS090562 to HSS, TE; R01 EY027036, R01 NS104926, UH2 CA203710 to HSS, R01 EY036978 to GS), the German Research Foundation (DFG; SPP 2041, 313856816 to PB, TE), the European Research Council (ERC, NextMechMod, 101039115 to PB), the Hertie Foundation (to PB), and through the Transformational Team Science Award from Research to Prevent Blindness (RPB; to GS, PB, TE, HSS).

## AUTHOR CONTRIBUTIONS

Conceptualization: SS, KF, GS, PB, HSS, TE;

Data curation: SS, SE, JF, JL, JO, DB, DG, CD, MS, NS, JS, YT, HM, NO, GS;

Formal analysis: SS, SE, JL, JO;

Investigation: SS, SE, JF, JL, JO, KF, DB;

Methodology: SS, SE, JF, JL, JO, KF;

Funding acquisition: GS, PB, HSS, TE;

Project administration: SE, JF, JL, CD, MS;

Resources: CD, MS, TS, PB, HSS, TE,;

Software: SS, JL, JO, ZH, RL, CD, AM, VS, GS;

Supervision: GS, PB, HSS, TE;

Validation: SS, SE, JF, JL, JO, DB, TS, GS;

Visualization: SS, SE, JF, JL, JO, AS, TE;

Writing – original draft: SS, SE, JL, JO, TE;

Writing – review & editing: SS, SE, JF, JL, JO, GS, PB, HSS, TE;

## Supplementary Note 1: Imaged EM stack

**Figure S1.**
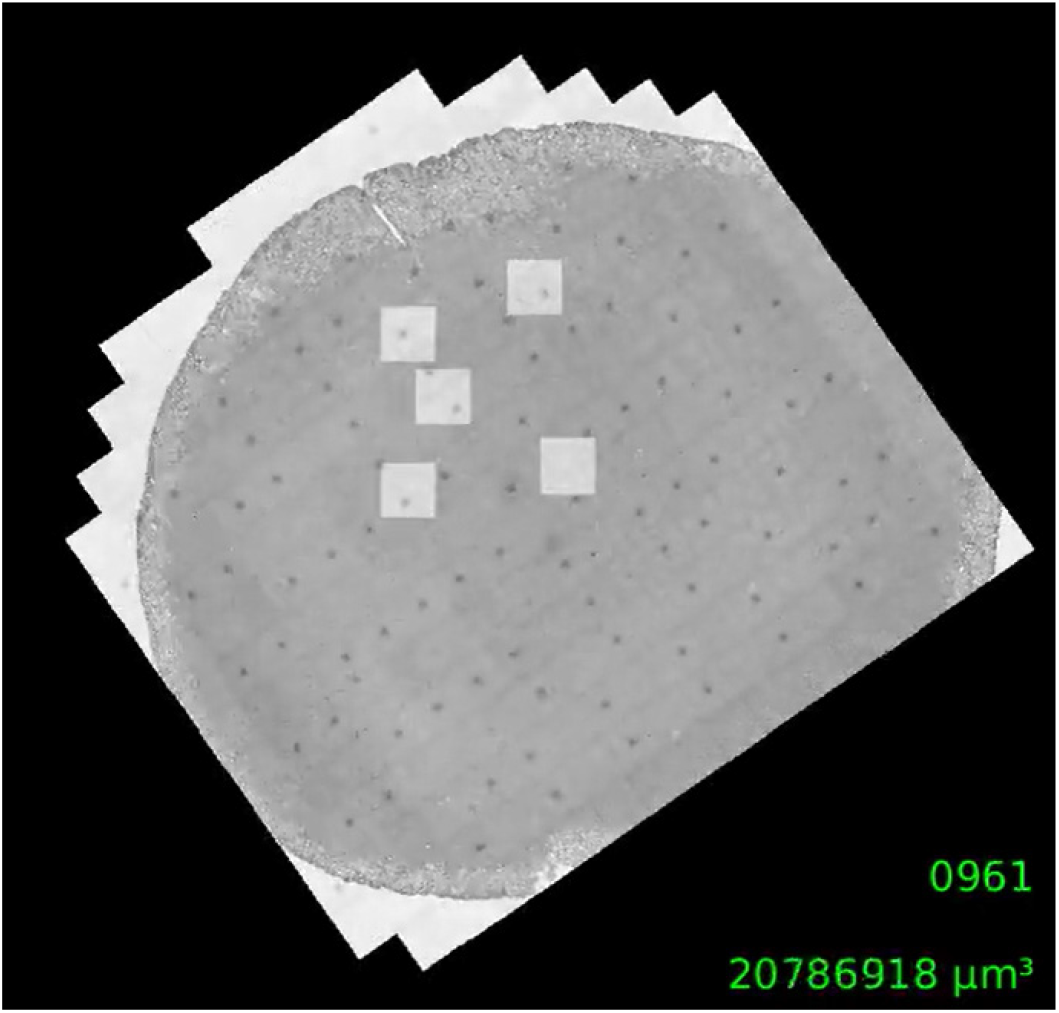
Supplement 1 to Fig. 1 – Movie of imaged EM stack. See Movie-figure1-supplemental-1.mp4.

## Supplementary Note 2: BC ribbon count and size distributions

**Figure S2.**
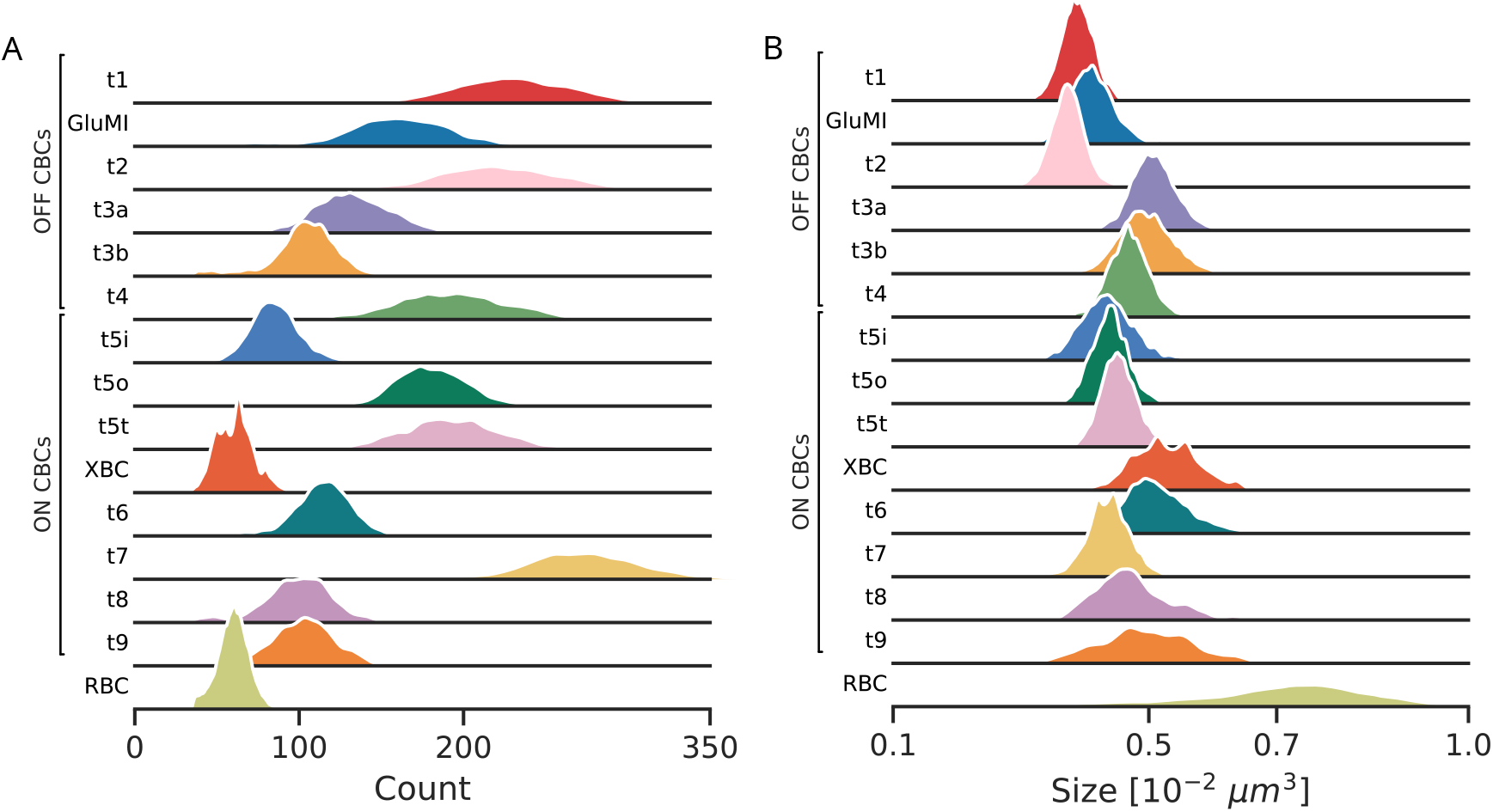
Supplement 1 to Fig. 8 – BC ribbon count and size distributions. (**A**) Normalized distributions of ribbon count per cell for each BC type. (**B**) Normalized distributions of mean ribbon size per cell for each BC type. Same dataset and BC type colors as in Fig. 8F and Fig. 4.

## Supplementary Note 3: BC ribbon statistics

**Figure S3.**
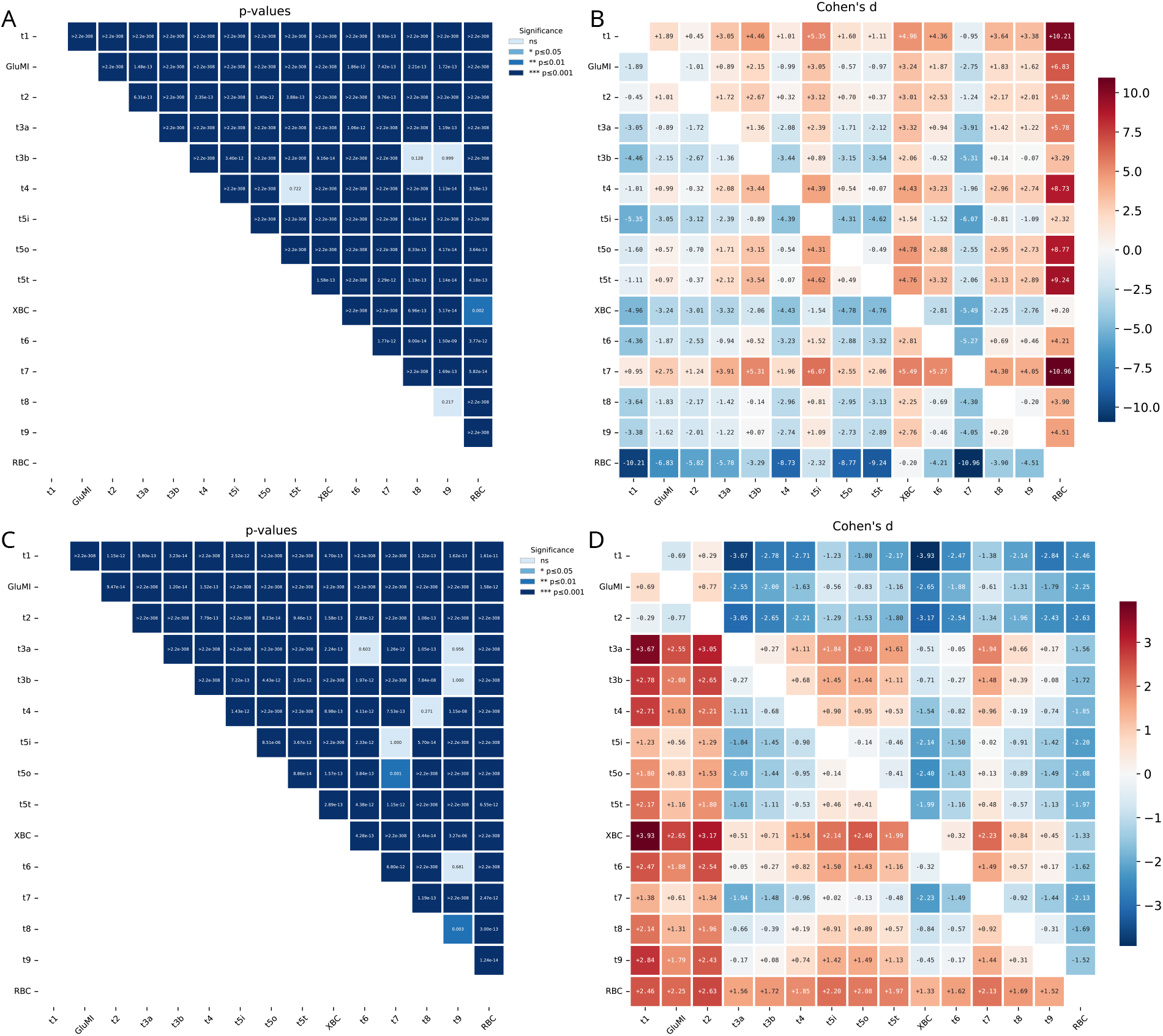
Supplement 2 to Fig. 8 – BC ribbon statistics. (**A**,**B**) p-values ((A), post-hoc pairwise Games-Howell test) and effect size ((B), Cohen’s *d*) for ribbon counts. (**C**,**D**) Same as in (A,B) but for mean ribbon size. Same dataset and BC type colors as in Fig. 8F and Fig. 4.

